# The WalRK two-component system in *Streptococcus pneumoniae* ensures robustness of secondary wall polymer attachment

**DOI:** 10.64898/2026.03.25.714115

**Authors:** Justin J. Zik, Zeyu Fu, Morgan N. Price, Yujie Li, Yuan Qiao, Adam P. Arkin, Adam M. Deutschbauer, Josue Flores-Kim, Lok-To Sham

## Abstract

Capsular polysaccharide (CPS) is essential for *Streptococcus pneumoniae* virulence. Yet, the mechanism linking CPS to peptidoglycan (PG) remains unclear. Here, we identified a strong negative genetic interaction between the genes encoding the putative capsule ligase CpsA and the WalK histidine kinase, a component of the WalRK two-component system regulating cell wall homeostasis. In the absence of *cpsA*, capsule polymers compete with wall teichoic acids for ligase activity to PG. This induces cell wall stress and is sensed by the WalRK system. Overexpression of the PG hydrolase *pcsB* or disruption of the PG-modifying enzymes *pgdA* and *oatA* restored growth of strains lacking *cpsA* and *walK*. Furthermore, CpsA overproduction compensates for the loss of other LytR-Cps2A-Psr (LCP) ligases, suggesting it can support capsule and wall teichoic acid syntheses. These findings support the model that LCP ligases are semi-redundant, although they may install secondary polymers on a different residue of PG. This work also suggests that WalRK signaling compensates for reduced capsule and WTA attachment by positively regulating PG hydrolases.

**SIGNIFICANT STATEMENT:** *Streptococcus pneumoniae* causes approximately half a million deaths annually. A powerful public health tool for controlling pneumococcal infections is vaccination against the protective capsule. Yet, the mechanisms by which the capsule layer attaches to the underlying cell wall remain poorly defined. This study shows that the conserved capsule gene CpsA is not strictly required for capsule attachment but instead works together with other LytRIZCpsAIZPsr (LCP) ligases. However, it requires the essential WalRK signaling system to maintain cell envelope integrity. Defects in LCP activity are alleviated by WalRKIZdriven upregulation of peptidoglycan hydrolases, overexpression of PcsB, or inactivating peptidoglycan modifications that limit hydrolysis. These findings reveal coordination among flux to capsule synthesis, secondary wall polymer attachment, and cell wall remodeling.

**TEASER:** The WalRK two-component system responds to cell envelope stress caused by reduced LCP ligase activity.

## INTRODUCTION

*Streptococcus pneumoniae* (pneumococcus) poses a significant threat to public health (1). It is one of the five bacterial pathogens responsible for more than half of all global deaths from infections (2). In 2019, pneumococcus was estimated to cause approximately 820,000 deaths worldwide, with an age-standardized mortality rate of 11.4% (2). It is also a leading cause of lower respiratory tract infections, resulting in roughly 97.9 million episodes annually (3), with many affected being children under the age of five (2–5). The increasing prevalence of antibiotic-resistant clinical isolates (6) has complicated the treatment of invasive pneumococcal diseases. Consequently, the World Health Organization listed pneumococcus as one of the twelve priority pathogens for antibiotic development. To reduce the burden of pneumococcal infections, many countries, including Singapore, have implemented vaccination programs for children and the elderly (9, 10). The capsular polysaccharide (CPS), a surface-exposed glycan layer that protects the cell against complement deposition and opsonophagocytosis, is the target of all licensed pneumococcal vaccines (11). Each pneumococcal strain typically produces a distinct CPS, and more than 100 serotypes have been identified to date (12, 13). The composition and structure of the CPS polymer are crucial determinants of pneumococcal interactions with host cells, including Kupffer cells (14, 15) and epithelial cells of the respiratory tract (16), thereby influencing colonization and virulence.

The Wzx/Wzy-dependent pathway is responsible for producing most pneumococcal CPS types (12, 13, 17). In this pathway, CPS precursors are sequentially added to the universal lipid carrier undecaprenyl phosphate (C55-P) until the repeating unit is formed (**Fig. 1**). The completed lipid-linked precursor is then translocated across the cytoplasmic membrane by the capsule transporter CpsJ (18–20), after which the glycan portion is polymerized by the CpsH polymerase (**Fig. 1**). The polymerized CPS is then ligated by a separate ligase protein to the peptidoglycan (PG) layer, which is reported to be the C6 position of the N-acetylglucosamine (GlcNAc) residue of PG (21). C55-P is recycled upon the completion of the CPS synthesis pathway. Since C55-P is also required for the synthesis of PG and teichoic acids, disruptions in capsule synthesis after the committed step can be lethal, likely due to sequestration of C55-P, which inhibits other essential cell envelope biosynthetic processes (20, 22). Growth can be restored by acquiring suppressor mutations that block initiation of the CPS pathway, such as disruption of the phosphoglycosyl transferase CpsE (20, 22) (**Fig. 1**). However, the resulting mutant is unencapsulated.

**Fig. 1.**
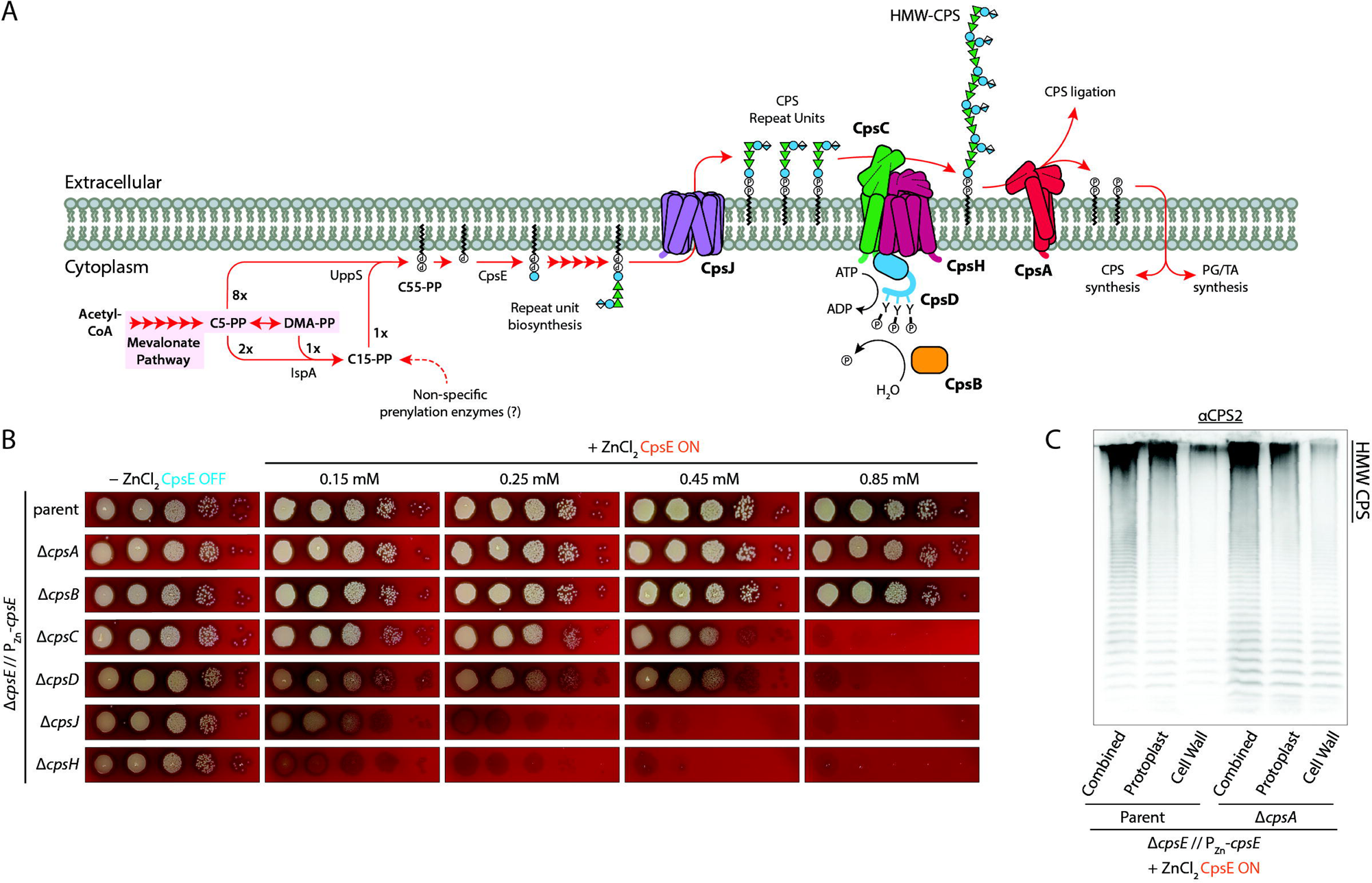
Blocking the completion of capsule synthesis is lethal. (**A**) Depicted is the serotype 2 capsule synthesis pathway of *S. pneumoniae*. Isopentyl pyrophosphate (C5-PP) is produced by the mevalonate pathway. It is the precursor of farnesyl pyrophosphate (C15-PP) that, together with the activity of UppS and additional C5-PP, generates undecaprenyl pyrophosphate (C55-PP). C55-PP is then dephosphorylated to make the lipid carrier C55-P. Capsule constituents are sequentially installed on C55-P until the capsule repeating unit is formed. The lipid-linked precursor is flipped by CpsJ, polymerized by CpsH, and attached to peptidoglycan. Biosynthesis of CPS releases C55-PP, which is then recycled for the synthesis of essential envelope polymers, such as wall teichoic acids. CpsBCD forms a tyrosine kinase system to control the length of the capsule polymers. CpsA is presumed to be the ligase, but supporting experimental evidence is lacking. (**B**) The inactivation of *cpsC*, *cpsD*, *cpsJ*, and *cpsH*, but not *cpsA* and *cpsB*, is lethal when CPS is produced. Strain NUS0267 [Δ*cpsE* // P_Zn_-*cpsE*] and its derivatives with clean deletions of the indicated genes were grown to the mid-exponential phase in BHI medium without the Zn^2+^ inducer, serially diluted, and spotted onto blood agar plates with the indicated concentration of Zn^2+^. Plates were incubated overnight at 37°C in 5% CO_2_ before imaging. The experiments were performed three times with similar results. (**C**) Cells without *cpsA* had a comparable amount of CPS attached to their surface. Strains NUS0267 and NUS0796 [Δ*cpsA* Δ*cpsE* // P_Zn_-*cpsE*] were grown in BHI supplemented with 0.45 mM of Zn^2+^ at 37°C in 5% CO_2_. Cultures were normalized by their optical densities and fractionated into cell wall and protoplast fractions. A combined fraction was also taken immediately prior to separation. Samples were treated with proteinase K and separated on 10% SDS-PAGE gels. CPS polymers were detected by immunoblotting using anti-serotype 2 antisera.

Except for serotype 37, genes required for CPS synthesis are organized into a single operon located between *dexB* and *aliA* (12, 17, 23). The first four genes of the capsule locus, *cpsA* (also known as *wzg*), *cpsB*, *cpsC*, and *cpsD* (collectively referred to as *wzbde*), are conserved (12, 17). *cpsA* encodes a putative capsule ligase that belongs to the LytR/Cps2A/Psr (LCP) family, whereas *cpsB, cpsC, and cpsD* form a tyrosine kinase system that regulates capsule polymer length (22). CpsC and CpsD are essential for the growth of the serotype 2 strain D39W on blood agar plates, likely because they help prevent C55-P sequestration by maintaining adequate capsule polymerase activity (22). They also recruit other capsule enzymes to the division septum (22, 24–27). Other serotype-specific genes, such as glycosyltransferases and the flippase, are located downstream of *cpsABCD* (12, 17, 23).

Although many factors involved in capsule synthesis in *S. pneumoniae* have been identified, the identity of the capsule ligase remains undetermined. LCP enzymes are implicated in ligating various secondary polymers to PG or a glycan acceptor (28–33), typically forming a phosphodiester linkage between them (29, 34, 35). However, biochemical experiments suggest that the connection between CPS and PG in pneumococcus is a direct glycosidic bond (21). This result challenges the role of CpsA as the capsule ligase because forming this type of linkage does not involve phosphoryl transfer (21). Contradictory evidence includes a report that hydrofluoric acid treatment can release the pneumococcal capsule, implying a phosphodiester bond between CPS and PG (26). Additionally, genetic inactivation of CpsA alone has yielded mixed results (36–38). Some mutants showed no detectable change in CPS attachment to PG (39, 40). One explanation is that LCP ligases are functionally redundant. Indeed, deletion of both *cpsA* and *lytR* results in the release of CPS into the medium, suggesting an inability to ligate CPS to PG (36, 37). Nevertheless, the Δ*cpsA* Δ*lytR* mutant exhibits increased cell lysis. Thus, it remains unclear whether the release is due to a lack of ligase activity (41).

In this study, we revisited the role of cpsABCD in capsule production. We confirmed that inactivation of *cpsA* does not significantly affect the length or proportion of CPS attached to PG. CpsD became dispensable when the interaction between CpsC and CpsH was maintained, suggesting that CpsD plays a major role as a scaffold protein. Randomly barcoded transposon sequencing (RB Tn-seq) experiments revealed a strong negative genetic interaction between *cpsA* and *walK*. This phenotype depended on capsule production and could be mitigated by overproducing the PG hydrolase PcsB. Consistently, deletion of *cpsA* led to *walK*-dependent upregulation of *pcsB*. Disruption of PG modifications by inactivating *pgdA* or *oatA* alleviated phenotypes associated with Δ*cpsA* Δ*walK*. Furthermore, we found that, when overexpressed, CpsA alone was sufficient to support growth in cells lacking all other LCP ligases, indicating that CpsA can ligate wall teichoic acid (WTA) polymers in pneumococcus. Finally, mutagenesis sequencing (Mut-seq) identified potential active-site residues in CpsA. Our results suggest a model in which, in the absence of CpsA, LytR and Psr link capsule polymers to PG, thereby reducing WTA ligation and PG remodeling. This defect could be corrected by deleting *pgdA* or *oatA*, or by overexpressing *pcsB* through the WalRK system. Our work clarifies the functions of conserved capsule genes and identifies a potential signal that activates the WalRK two-component regulatory system.

## RESULTS

### Disrupting capsule synthesis after the committed step is lethal

To examine the phenotypes of cells lacking late capsule synthesis genes, we constructed a strain in which the expression of the first enzyme in the pathway, CpsE, is under the control of a Zn^2+^-inducible promoter (Δ*cpsE* // P_Zn_-*cpsE*) (18). Depletion of *cpsE* allows late capsule enzymes to be inactivated without fitness defects by reducing flux through the pathway. When Zn^2+^ was added to the medium, cells lacking a functional *cpsC*, *cpsD*, *cpsJ*, or *cpsH* were unable to grow (**Fig. 1**). In contrast, the Δ*cpsA* and Δ*cpsB* strains remained viable. Δ*cpsA* did not lead to any detectable changes in CPS attachment, polymer length, or the proportion bound to peptidoglycan, and roughly equivalent amounts were found associated with the membrane and cell wall fractions after protoplasting, consistent with a previous report (**Fig. 1**) (39). CpsCD is an activator of CpsH (22). In its absence, cells accumulate partially polymerized, low-molecular-weight capsule polymers, which deprive the essential C55-P from other cell surface biogenesis pathways (22). In contrast, deletions of the *cpsJ* flippase and the *cpsH* polymerase likely lead to a more severe sequestration of C55-P via accumulation of C55-P linked to individual repeating units (20, 22). In line with their roles in capsule synthesis, we found that a stronger induction of *cpsE* is required to kill Δ*cpsC* and Δ*cpsD* strains compared to Δ*cpsJ* and Δ*cpsH* cells (**Fig. 1**).

### The requirement of CpsD for growth can be bypassed by bridging CpsC and CpsH

Many bacteria encode polysaccharide co-polymerases (PCPs) to regulate the activity of Wzy polymerases (42–44). Class 1 PCPs (PCP1), which include Wzz that modulates O-antigen synthesis, consist of two transmembrane helices and a large extracellular co-polymerase domain (42–44). Class 2a PCPs (PCP2a) have an additional tyrosine kinase domain at the C-terminus, comprising a Walker box and a YGX motif (42–44). This kinase domain is thought to autophosphorylate the tyrosine residues at the YGX motif. PCP1 and PCP2a are typically found in Gram-negative bacteria. In *S. pneumoniae*, CpsC and CpsD are part of the class 2b PCPs (PCP2b), where the tyrosine kinase domain (CpsD) is on a separate polypeptide chain from the co-polymerase (CpsC) (**Fig. 2**). Autophosphorylation of CpsD fine-tunes the co-polymerase activity of CpsC (22). This activity likely regulates the length of capsule polymers in various host niches (22). Although the kinase activity of CpsD is dispensable for growth, the CpsD protein itself is conditionally essential (22, 24). Δ*cpsD* cells fail to recruit other capsule enzymes to the divisome (24), suggesting that CpsD may also facilitate the interaction between CpsC and CpsH.

**Fig. 2.**
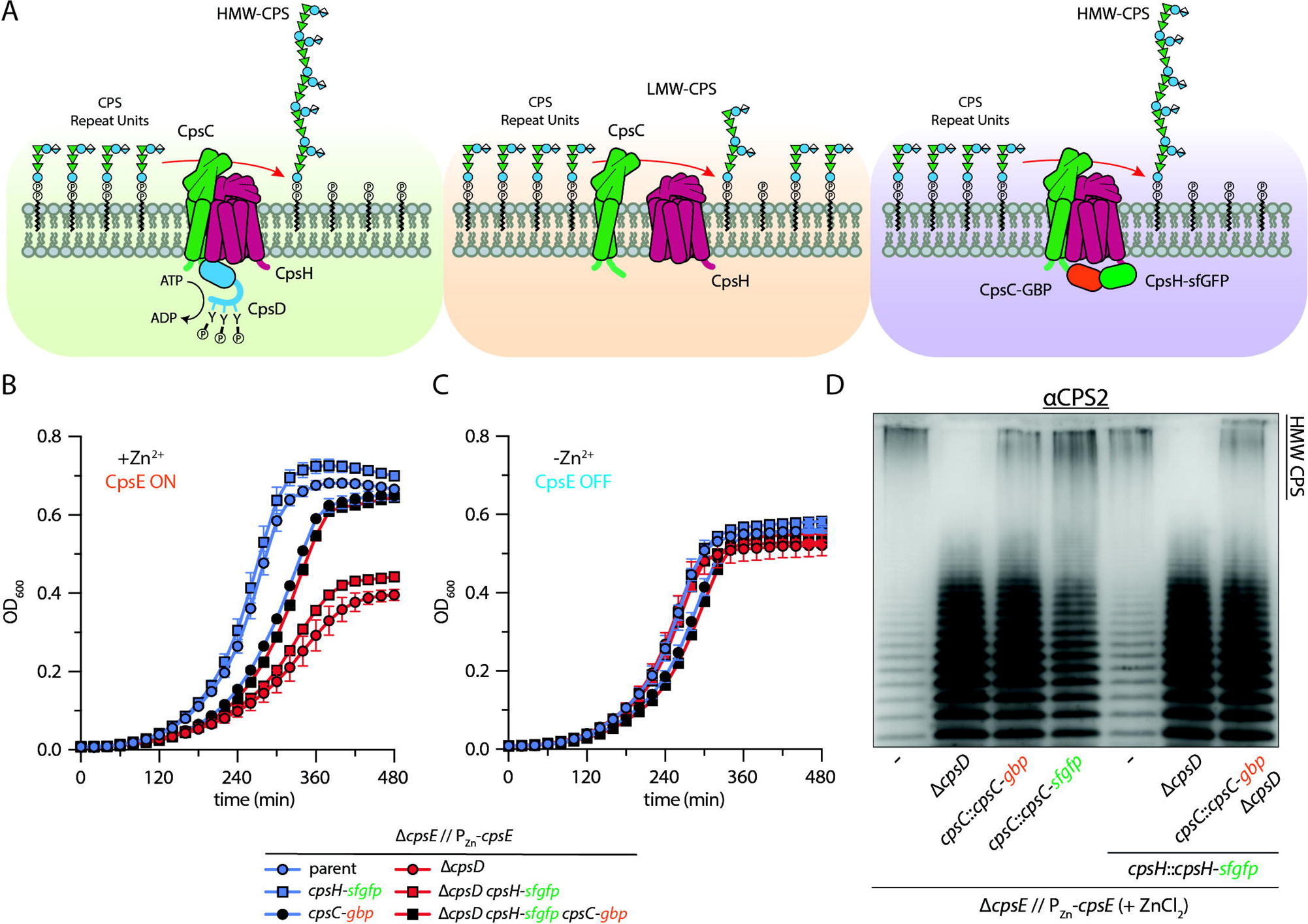
Bridging the interaction between CpsC and CpsH bypasses the requirement of CpsD for growth. (**A**) CpsC is a class 2b polymerase co-polymerase (PCP2b). Its activity is controlled by CpsD, which is an autokinase that phosphorylates its C-terminal tyrosine residues (green panel). Phosphorylation is thought to reduce the co-polymerase activity of CpsC, resulting in shorter capsule polymers. The inactivation of CpsD results in a drastic reduction in CpsH activity. When the capsule is produced, Δ*cpsD* leads to a sequestration of Und-P and stalls growth (orange panel). Forcing CpsC and CpsH to interact by fusing them with green fluorescence protein (GFP) and GFP-binding protein (GBP), respectively, bypasses the requirement of CpsD (purple panel). Strains NUS0267 [Δ*cpsE* // P_Zn_-*cpsE*] and NUS0680 [Δ*cpsD* Δ*cpsE* // P_Zn_-*cpsE*], and their derivatives expressing the indicated variants of CpsC and CpsH, were grown in BHI in the absence of Zn^2+^. Cultures were diluted in fresh BHI medium with (**B**) or without (**C**) 0.85 mM Zn^2+^ supplementation. Growth was monitored by measuring the optical densities of the culture over time. Shown as the error bars are the standard deviations from three biological repeats. (**D**) Artificially recruiting CpsC to CpsH partially restores high molecular weight (HMW) polymer production in the Δ*cpsD* mutant. Strains described in (**C**) were grown in BHI with 0.45 mM Zn^2+^ for one hour. Cells were harvested and resolved on an SDS-PAGE gel. CPS was detected by immunoblotting using the anti-serotype 2 (anti-CPS) antibodies. Experiments were performed three times with similar results.

To test this hypothesis, we fused the C-terminus of CpsH with green fluorescent protein (GFP) and CpsC with a GFP-binding protein (GBP) (45). Cell viability was maintained by regulating *cpsE* expression (Δ*cpsE* // P_Zn_-*cpsE*). Both constructs remained partially functional, as evidenced by the formation of high-molecular-weight (HMW) CPS polymers, but with mild growth defects and accumulation of low-molecular-weight CPS (**Fig. 2**). *cpsD* became dispensable for growth only when both CpsC and CpsD were tagged by GFP and GBP, respectively (**Fig. 2**). The resulting Δ*cpsD* strain produced HMW CPS polymers similar to the isogenic *cpsD*^+^ strains expressing one of the fusion constructs (**Fig. 2**). Our results indicate that bridging the interaction between CpsC and CpsH is sufficient to bypass the requirement of CpsD for growth on blood agar plates. In wild-type cells, when CpsD is absent, CpsC cannot adequately activate CpsH, resulting in the lethal sequestration of C55-P (22).

### RB Tnseq screens reveal genetic interactions of CPS genes

Next, we performed random barcode transposon sequencing (RB Tn-seq) (46) to identify genetic interactions in cells lacking either *cpsA*, *cpsB*, *cpsC*, or *cpsD* when the capsule is produced. To do this, we first grew the *cpsE*-inducible strain (Δ*cpsE* // P_Zn_-*cpsE*) and its derivatives with either Δ*cpsA*, Δ*cpsB*, Δ*cpsC*, or Δ*cpsD* in the absence of Zn^2+^ and transformed each with genomic DNA from a previously mapped RB Tn-seq library (47). We then plated these transformants on media supplemented with a sublethal level of Zn^2+^ inducer (0.45 mM). After overnight incubation, we collected the surviving cells and extracted their DNA for barcode sequencing. As predicted from the lack of phenotype of the Δ*cpsB* mutant, we did not identify strong genetic interactions associated with this gene (**Fig. S1, table S3**). By contrast, several significant negative genetic interactions were identified in the RB Tn-seq screen of strain NUS0679 (Δ*cpsC* Δ*cpsE* // P_Zn_*-cpsE*) (table S3). Some of these genes are associated with C55-P or cell envelope synthesis, such as *ispA*, *panT*, and *tacL* (**Fig. 3**). IspA is a farnesyl diphosphate synthase that produces a precursor of C55-P (i.e., farnesyl diphosphate) from two isopentyl diphosphate (IPP) and one dimethylallyl diphosphate (DMAPP) molecules (48) (**Fig. 1**). PanT is a putative substrate-specific component of an energy-coupling factor (ECF) transporter that is specific to pantothenate. Its loss likely impairs acetyl-CoA synthesis and indirectly inhibits the mevalonate pathway and C55-P synthesis (49). TacL is a lipoteichoic acid (LTA) ligase that attaches the teichoic acid precursor to glucosyl diacylglycerol (34). A deficiency in TacL results in an inability to regulate the autolysin LytA (50). Thus, Δ*tacL* may exacerbate phenotypes associated with mutations that reduce PG synthesis. Similar genetic interactions were also identified in a parallel screen performed in NUS0680 (Δ*cpsD* Δ*cpsE* // P_Zn_*-cpsE*), likely because of the overlapping function of CpsC and CpsD (**Fig. S1**). To validate the RB Tnseq results, we deleted these genes in the NUS0679 (Δ*cpsC* Δ*cpsE* // P_Zn_*-cpsE*) background and demonstrated that the synthetic lethality/sickness phenotypes were reproducible (**Fig. 3**). These results align with the model that losing *cpsC* or *cpsD* causes C55-P sequestration, which leads to cell envelope stress and decreased viability.

**Fig. 3.**
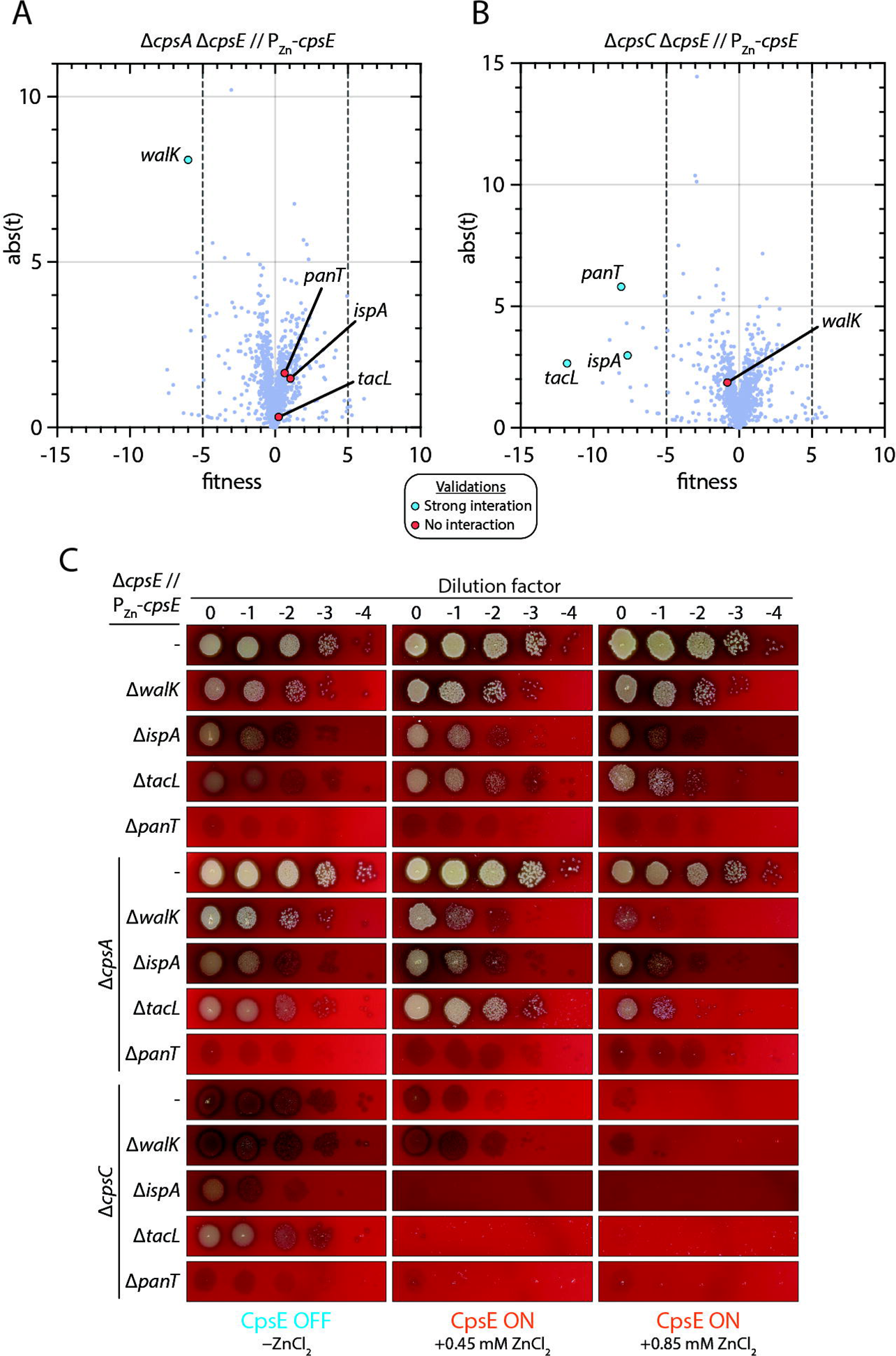
*cpsA*, but not *cpsC*, exhibits a negative genetic interaction with *walK*. RB Tn-seq uncovered genetic interactions of *cpsA* (**A**) and *cpsC* (**B**). Strains NUS0796 [Δ*cpsA* Δ*cpsE* // P_Zn_-*cpsE*] and NUS0679 [Δ*cpsC* Δ*cpsE* // P_Zn_-*cpsE*] were grown in BHI without Zn^2+^, transformed with a barcoded Tn library, and plated on blood agar supplemented with 0.45 mM Zn^2+^. Cells on the plates were scraped and pooled, DNA was extracted, and the barcodes were amplified and sequenced. Blue dots represent genetic interactions that were reproduced when the corresponding strains were reconstructed. Red dots indicate deletions that did not exhibit detectable genetic interactions. (**C**) Spot dilution assays showing the genetic interactions involving *cpsA* and *cpsC*. Strains NUS0267 [Δ*cpsE* // P_Zn_-*cpsE*], NUS0679, NUS0796, and their derivatives that lack the indicated genes were grown in BHI until the exponential phase in the absence of Zn^2+^. Cultures were serially diluted and spotted on blood agar plates with or without Zn^2+^ supplementation. Plates were incubated overnight at 37°C in 5% CO_2_ before imaging. Experiments were performed three times with similar results.

Compared to CpsBCD, the role of CpsA in capsule synthesis is less clear. CpsA is recruited to the septum by CpsC (24, 25, 36). Mislocalization of CpsA by inactivating CpsC (24), or by introducing a mutation in CpsC (V56A) (25), results in a reduction of capsule materials at *S. pneumoniae* midcell, consistent with the hypothesis that CpsA is responsible for decorating the cell surface with capsule polymers. Nevertheless, as in a previous report (39), we found that Δ*cpsA* did not appear to affect CPS attachment to peptidoglycan (**Fig. 1**). This suggests that other enzymes may compensate for capsule ligase activity in pneumococcus. To test this hypothesis, we performed RB Tn-seq on strain NUS0796 (Δ*cpsA* Δ*cpsE* // P_Zn_-*cpsE*) in the presence or absence of the Zn^2+^ inducer. This screen is expected to report gene(s) that are synthetically lethal or sick with *cpsA*, assuming that capsule ligase activity is required for growth, analogous to other late-stage capsule enzymes. Unexpectedly, we did not observe negative genetic interactions between *cpsA* and *panT*, *ispA*, or *tacL* in this screen, suggesting that cells lacking *cpsA* do not severely sequester C55-P. Instead, we found *cpsA* has a strong genetic interaction with the histidine kinase *walK*. Cells lacking *cpsA* and *walK* formed chains and occasionally bulged (**Fig. 4**). Inducing capsule production in these cells (NUS3698 [Δ*walK* Δ*cpsA* Δ*cpsE* // P_Zn_-*cpsE*]) seemed to reduce bulging, likely because CPS masks cell wall defects (51). However, they became phase-bright and could not form colonies on blood agar plates (**Fig. 4**). The phase-bright cells were probably not ghosts, as they were not exclusively labeled with propidium iodide (**Fig. S2**). The basis for this phase-bright appearance is unclear. A similar phenomenon has been reported in Bacillus spp. endospores, where phase-brightness reflects an increased refractive index caused by core dehydration and the accumulation of calcium dipicolinate (Ca-DPA), which together raise the optical density of the spore interior (52). In *S. pneumoniae*, loss of WTA attachment (Δ*lytR*) reduces cell wall thickness by ∼36% (53), possibly lowering the overall refractive index of the cell periphery. In liquid growth, the Δ*cpsC* Δ*walK* mutant was also sick. Nevertheless, this appeared to be the combined effect of growth defects caused by Δ*walK* and Δ*cpsC* when capsule production was induced, and it was not as pronounced as the Δ*cpsC* Δ*ispA* double mutant (**Fig. S3**). To eliminate the possibility of polar effects, we demonstrated that these phenotypes could be rescued by expressing *cpsA* and *walRKJ* ectopically (**Fig. 4**). The observation that the Δ*cpsA* Δ*walK* mutant was unable to form colonies when *cpsE* was expressed indicates that this negative genetic interaction requires capsule synthesis, but is unlikely to be caused by C55-P sequestration.

**Fig. 4.**
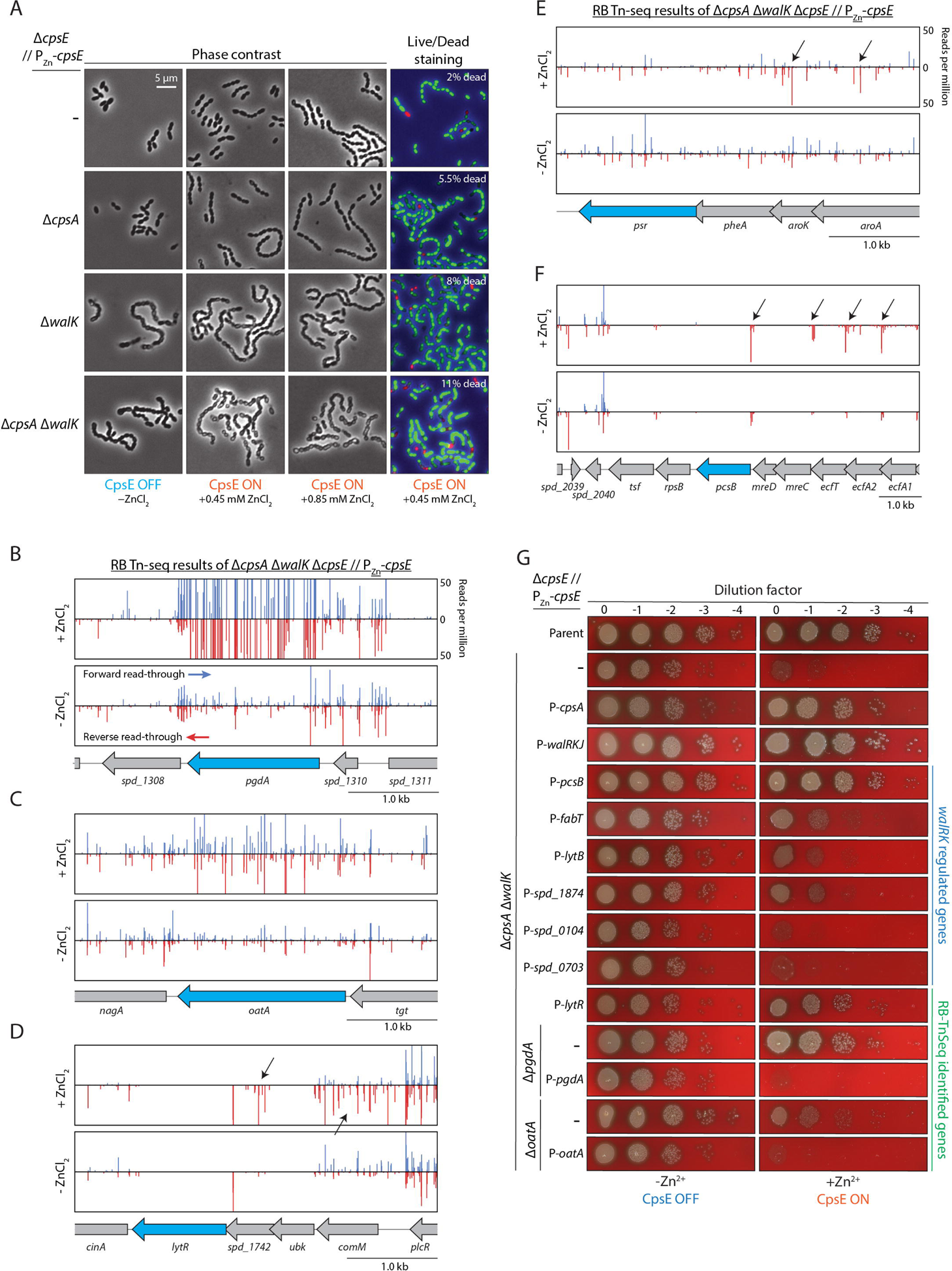
The synthetic lethality of Δ*cpsA* and Δ*walK* can be bypassed by overexpressing *pcsB* or *lytR*, or by deleting *pgdA* or *oatA*. (**A**) Cells deprived of *walK* and *cpsA* exhibited morphological defects when capsule production was induced. Strain NUS0267 [Δ*cpsE* // P_Zn_-*cpsE*] and its derivatives lacking *cpsA* (NUS0796), *walK* (NUS3688), and both (NUS3698) were grown in BHI without Zn^2+^, normalized by optical density, and then released into BHI with or without 0.85 mM Zn^2+^. After five doublings, cells were collected and visualized by phase-contrast microscopy. Alternatively, they were stained with the LIVE/DEAD BacLight Bacterial Viability Kit (Molecular Probes) to assess cell viability. Cells stained with SYTO 9 appear green (live), while cells stained with propidium iodide appear red (dead). Images were processed using Fiji (101). The brightness and contrast of each channel were adjusted in a similar manner to facilitate visual comparison between the two populations. Next, to identify conditions that bypass the lethality caused by Δ*walK* Δ*cpsA*, strain NUS3698 [Δ*cpsA* Δ*walK* Δ*cpsE* // P_Zn_-*cpsE*] was grown in BHI, transformed with a barcoded Tn library, and plated in the absence of Zn^2+^. Transformants were collected and subsequently plated again in the absence or presence of Zn^2+^. Shown are the RB Tn-seq results around the genetic loci of *pgdA* (**B**), *oatA* (**C**), *lytR* (**D**), *psr* (**E**), and *pcsB* (**F**). An increase in transposon insertions in a single orientation, which may have led to overexpression of downstream genes, is indicated by the black arrows. (**G**) Validation of the RB Tn-seq results. Strain NUS0267 [Δ*cpsE* // P_Zn_-*cpsE*] and its derivatives with the indicated modifications were grown in BHI without the Zn^2+^ inducer at 37 °C in 5% CO_2_. Cultures were serially diluted and spotted onto blood agar plates with or without Zn^2+^, incubated overnight at 37 °C in 5% CO_2,_ and imaged. Experiments were conducted three times with similar results.

### The genetic interaction between *cpsA* and *walK* is linked to defective PG remodeling

The WalRK system is widely conserved in Firmicutes (reviewed in (54–57)). In general, WalRK regulates genes in response to cell envelope stresses. Phosphorylation of the WalR response regulator increases the expression of PG hydrolases, as reported in many species, including *Bacillus subtilis* (58), *Staphylococcus aureus* (59), and *S. pneumoniae* (60). In pneumococcus, it is the only two-component system (TCS) required for growth under laboratory conditions (61–63). Additionally, only the WalR response regulator is essential, while the WalK histidine kinase is dispensable (62). The requirement of the WalRK TCS for growth is often attributed to the essentiality of the cell wall hydrolases it regulates, since constitutive expression of the essential hydrolase gene *pcsB* is sufficient to render *walR* non-essential (61–63). Although the exact signals sensed by the WalK histidine kinase are not fully understood, it is thought to detect intermediates related to cell envelope synthesis (54–57). In *B. subtilis*, the extracellular sCache domain of WalK detects PG fragments released by LytE and CwlO, thereby regulating their expression levels (58). However, the sCache domain is absent in the pneumococcal WalK (55). Unlike in *B. subtilis*, WalK homologs in Streptococci are not essential for growth (e.g., *S. pneumoniae*, *Streptococcus mutans*, *Streptococcus pyogenes*) (62, 64, 65). Since the extracellular domain of the pneumococcal WalK is unusually small, it likely senses a yet-to-be-identified cytoplasmic signal, possibly through its conserved Per-ARNT-Sim (PAS) domain (58, 66, 67).

The requirement of the WalRK system for growth can be bypassed by constitutive expression of a PG hydrolase, PcsB, which has recently been shown to be a D,L-endopeptidase (62, 68). Thus, if the Δ*walK* Δ*cpsA* lethality is caused by misregulation of the WalRK regulon, compensating for its function by ectopically expressing *pcsB* should rescue the phenotype. Indeed, growth was restored when the constitutive P-*pcsB* cassette was introduced into the Δ*walK* Δ*cpsA* mutant (**Fig. 4**). Additionally, we observed a partial rescue of the phenotype when other WalR-regulated genes were constitutively expressed (P-*fabT*, P-*lytB*, and P-*spd_1874*). As LytB and Spd_1874 (or VldE) are also PG hydrolases (69, 70), they may partially substitute for PcsB functions. Why the constitutive expression of FabT alleviates the Δ*walK* Δ*cpsA* phenotype remains unclear. Since it controls many genes involved in fatty acid synthesis (71), it may indirectly affect PG or wall teichoic acid (WTA) synthesis, for example, by altering cellular levels of diacylglycerol or Und-P.

To understand the genetic interaction between Δ*walK* Δ*cpsA*, we performed RB TnSeq on strain NUS3698 in the presence or absence of Zn^2+^ inducer. Tn insertions that improved the growth of the Δ*walK* Δ*cpsA* mutant should be enriched under non-permissive conditions (i.e., when capsule expression was induced with Zn^2+^ supplementation). Conversely, insertions that exacerbated the Δ*walK* Δ*cpsA* phenotype would be less frequent. As expected, we saw an increase in Tn insertions upstream of *pcsB* with the Tn-encoded promoter oriented in the same direction as the open reading frame (**Fig. 4**). Because the Tn lacks a transcriptional terminator, insertions upstream of *pcsB* likely increase its expression via read-through transcription originating from the constitutive promoter that drives the antibiotic resistance cassette (47). Similarly, more Tn insertions were found upstream of *lytR*, suggesting possible interchangeability of the LCP enzymes. Additionally, Tn insertions presumed to disrupt the function of *pgdA* and *oatA* were also enriched when capsule production was induced, indicating that the absence of PG-modifying enzymes alleviates the Δ*walK* Δ*cpsA* phenotype. We validated these results by reconstructing the strains in the NUS3698 background to confirm the phenotype (**Fig. 4**). Expectedly, complementing *pgdA* and *oatA* restored the lethality of the Δ*walK* Δ*cpsA*in the Δ*pgdA* and Δ*oatA* mutants, respectively, when capsule expression is induced (**Fig. 4**).

One reason WalK may be required for growth in Δ*cpsA* cells is that it detects and responds to specific cell envelope defects resulting from disruptions in the capsule pathway. Consistent with this hypothesis, *pcsB* expression increased by about fourfold in the Δ*cpsA* mutant upon expression of *cpsE* (**Fig. 5**). This increase depended on capsule production and the presence of *walK* (**Fig. 5**). Next, we examined whether the requirement of WalK for viability is specific to the Δ*cpsA* mutant or if it is a general requirement when LCP protein activity is reduced. To test this, we constructed a strain in which *cpsE* was deleted, and *lytR* was under a zinc-inducible promoter. Under these conditions, cells depleted of *lytR* could form colonies on blood agar plates, but not if *walK* was also inactivated (**Fig. 5**). These results suggest that *walK* becomes essential when the cell lacks LCP activity, likely because it positively regulates *pcsB* expression to overcome WTA stress on the cell surface (**Fig. 4**).

**Fig. 5.**
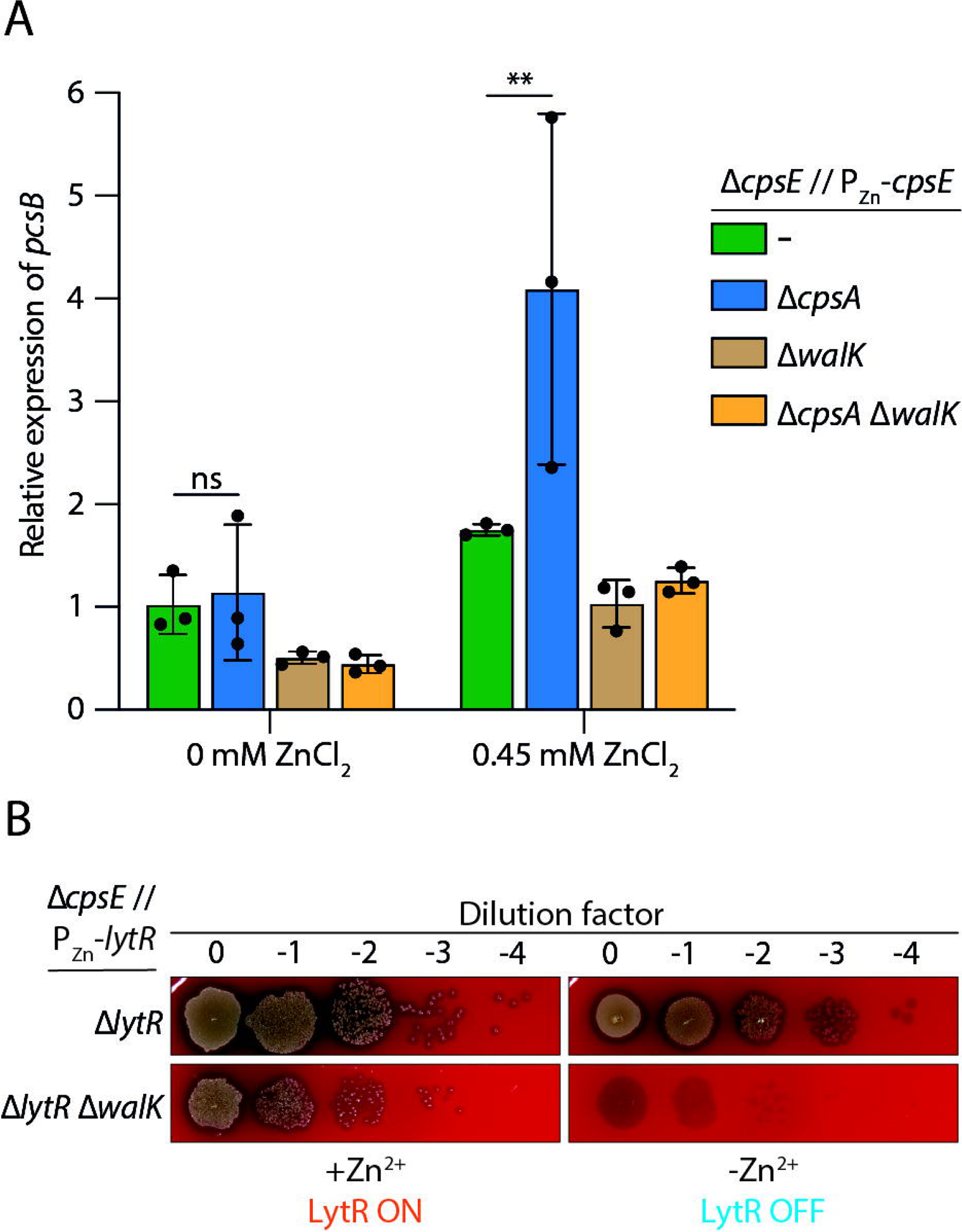
Inactivating *cpsA* increases *pcsB* expression through WalRK signaling. (**A**) Strains NUS0267, NUS0796, NUS3688, and NUS3698 were initially grown in BHI at 37 °C in 5% CO_2_ without Zn^2+^, then released into 0.45 mM Zn^2+^ and incubated as above for one hour. Cells were mixed with RNAprotect Bacteria Reagent (Qiagen) and harvested by centrifugation. RNA was extracted, and the relative amount of the pcsB *transcript* was measured by quantitative reverse transcription PCR (qRT-PCR). Plotted are the means and standard deviations from three biological repeats. P-values were calculated by two-way ANOVA followed by Tukey’s multiple comparisons test. ns, not significant; **, p < 0.01. (**B**) WalK is required for growth in cells lacking LytR. Strains NUS5965 [Δ*cpsE* Δ*lytR* // P_Zn_-*lytR*] and NUS7076 [Δ*cpsE* Δ*lytR* Δ*walK* // P_Zn_-*lytR*] were grown in BHI at 37 °C in 5% CO_2_ supplemented with 0.45 mM Zn^2+^. When the cultures reached the exponential phase, cells were serially diluted and spotted on blood agar plates with or without 0.45 mM Zn^2+^. Plates were incubated overnight at 37 °C in 5% CO_2_ before imaging. Shown are the representative images from three biological repeats.

Next, we tested whether increasing the amount of WTA on the cell surface could restore growth of the Δ*cpsA* Δ*walK* mutant. In *S. pneumoniae*, TacL is the lipoteichoic acid (LTA) ligase that attaches the teichoic acid component to glucosyl-diacylglycerol (Glc-DAG) (34, 50) (**Fig. S5**). Since LTA is not required for growth under lab conditions (34, 50), the loss of TacL is known to redirect more teichoic acid precursors to the WTA pathway (50). We thus constructed Δ*tacL* mutants in the NUS0267 [Δ*cpsE* // P_Zn_-*cpsE*] background, with or without Δ*cpsA*, Δ*walK*, or both, and tested whether inducing capsule production was tolerated. Under this condition, cells lacking *tacL* did not significantly restore growth of the Δ*cpsA* Δ*walK* mutant (**Fig. S5**). Another way to increase WTA levels on the cell surface is to disrupt the WTA hydrolase WhyD (72). In the Δ*whyD* background, capsule production appeared lethal even in the *cpsA*^+^ *walK*^+^ parent (**Fig. S5**). Introducing the Δ*cpsA* cassette further exacerbated the Δ*whyD* phenotype, but only in a capsule-production-dependent manner (**Fig. S5**). Thus, the growth defects caused by Δ*cpsA* Δ*walK* mutations could not be explained solely by a reduction in WTA on the cell surface.

### Inactivating PG-modifying enzymes alleviates the chaining phenotype of Δ*walK* strains

Δ*walK* cells form chains (67). This phenotype can be rescued by increasing *pcsB* expression (51, 73). Since Δ*pgdA* and Δ*oatA* reduce the synthetic sickness caused by Δ*walK* Δ*cpsA*, we wondered if they could also correct the chaining phenotype. To investigate this, we quantified the proportion of chaining cells by measuring side (SSC-A) and forward scattering (FSC-A) using flow cytometry (**Fig. S4**). Cells lacking CpsA were indistinguishable from the wild-type in terms of the proportion of cells that form chains (**Figs. 6 and S6**), indicating that the absence of WalK primarily caused the chaining phenotype. Combining Δ*cpsA* and Δ*walK* did not worsen the chaining phenotype (**Fig. 6**). Complementing *walK* by introducing the P-*walRKJ* cassette corrected the chaining in the Δ*cpsA* Δ*walK* mutant (**Fig. 6**). Similarly, ectopic expression of *lytR* also partially reduced chaining, suggesting that an increase in LCP activity may alleviate Δ*walK* defects, possibly by increasing the activity of PG hydrolases. Expectedly, expressing *pcsB in trans* fully restored chaining in Δ*cpsA* Δ*walK* cells (**Fig. 6**). *lytR* and *cpsA* may be partially interchangeable because introducing the P-*lytR* cassette corrected the chaining phenotype in the Δ*walK* mutant, but to a lesser extent in the Δ*cpsA* Δ*walK* double mutant (**Fig. 6**). This result was unlikely due to the role of *cpsA* in the capsule pathway because *cpsE* was not induced. Contrary to their ability to restore the growth of the Δ*cpsA* Δ*walK* mutant (**Fig. 4**), inactivation of *oatA*, but not *pgdA*, reduced chaining in the Δ*walK* mutant (**Fig. 6**). Complementing *oatA* restored chaining. These results show that reducing PG modifications by deleting enzymes responsible for O-acetylation and N-deacetylation may increase the PG hydrolase activity required for splitting daughter cells.

**Fig. 6.**
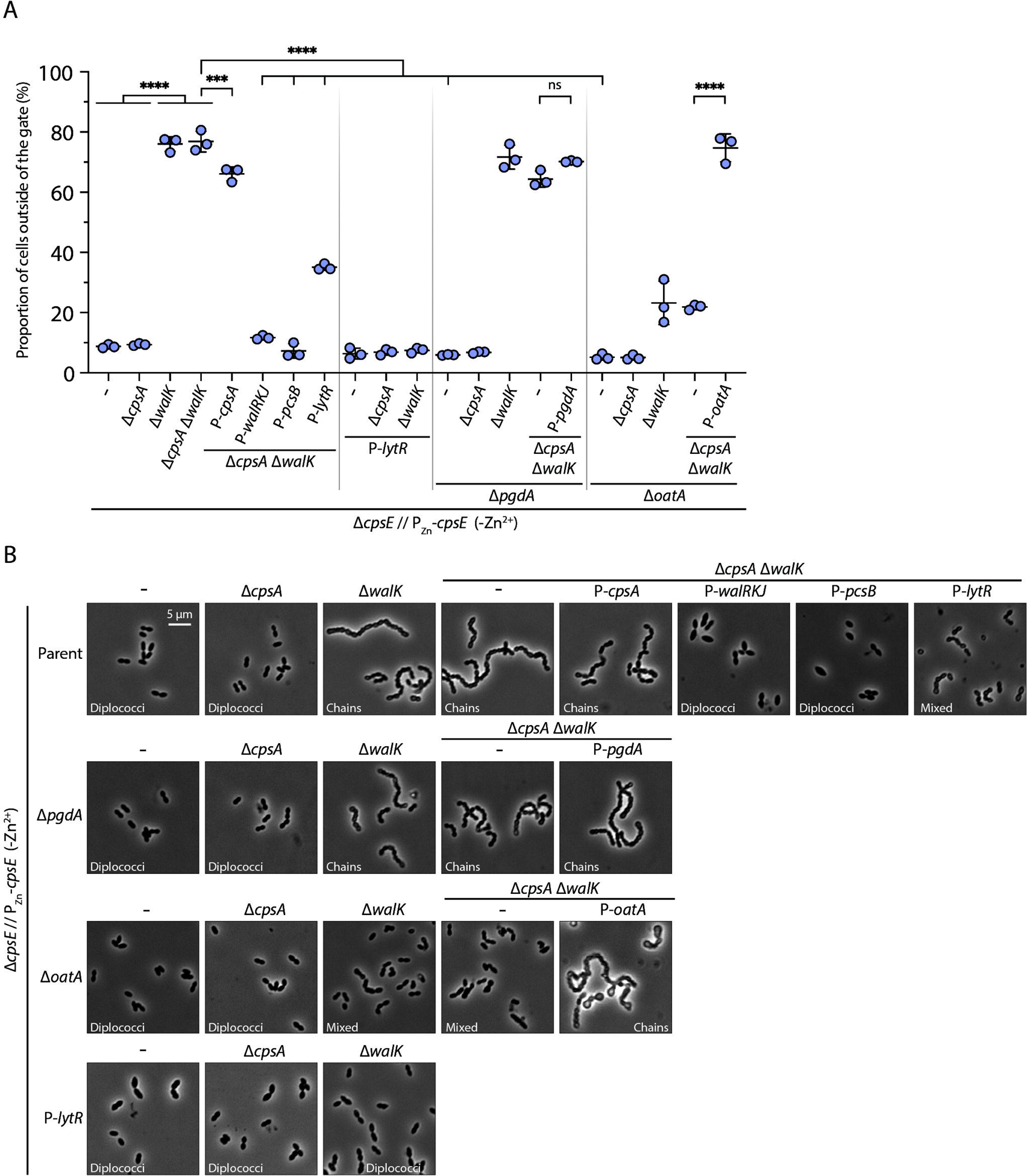
Δ*oatA*, but not Δ*pgdA*, alleviates the chaining phenotype of Δ*walK*. (**A**) Flow cytometry experiments were conducted to evaluate the chaining phenotype in strains lacking *walK* and *cpsA*. Strain NUS0267 [Δ*cpsE* // P_Zn_-*cpsE*] and its derivatives with the indicated modifications were grown in BHI without the Zn^2+^ inducer at 37 °C in 5% CO_2_ until early log phase, inactivated with 1% paraformaldehyde, and analyzed on a flow cytometer. Cells that form chains cause an increase in side-scattering (SSC) and forward-scattering (FSC) and therefore fall outside of the gate where about 90% of the parent strain falls (**Fig. S6**). Plotted are the means and standard deviations from three biological repeats. P-values were calculated by one-way ANOVA followed by Tukey’s multiple comparisons test. ****, p < 0.0001; ***, p < 0.001. (**B**) Phase contrast microscopy of the cells from strains described in (**A**) under the same growth conditions. Bar, 5 µm. Shown are the representative micrographs from three biological repeats.

### CpsA, LytR, and Psr are non-redundant but interchangeable when overexpressed

Our genetic results suggest the following model: When *cpsA* is deleted, LytR and Psr attach capsule polymers to PG. Since WTA polymers are primarily linked to PG by LytR and Psr, additional capsule polymers may overload these ligases, resulting in a deficiency of WTA on the cell surface. The WalRK TCS detects this shortage and triggers a response to mitigate this cell wall stress, such as by increasing *pcsB* expression. Why higher PcsB levels suppress the Δ*cpsA* phenotype remains unclear. The inability to ligate WTA may lead to the accumulation of lipid-linked polymers on the outer leaflet of the cell membrane. Possibly, these polymers decrease PcsB activity, similar to a reduction in overall PG hydrolase activity in the LCP mutants of *Bacillus subtilis* (74).

Consistent with this hypothesis, we were unable to construct the Δ*cpsA* Δ*lytR* double mutant in the *cps*^+^ background. Therefore, we inactivated *cpsE* and inserted the P_Zn_-*lytR* cassette at an ectopic site (NUS5935 [Δ*cpsE* // P_Zn_-*lytR*]). Next, we inactivated various LCP proteins singly or in combination. Even when *lytR* was depleted, Δ*cpsA* and Δ*psr* mutants could still form colonies on blood agar plates (**Fig. 7**). However, if *cpsA* and *psr* were simultaneously inactivated, cells could not grow unless *lytR* expression was restored (**Fig. 7**). These phenotypes were more prominent in BHI, as Δ*cpsA* alone generally exacerbated the growth defects caused by Δ*lytR* or Δ*lytR* Δ*psr* (**Fig. 7**). Cells lacking LytR and other LCP ligases form chains, and were occasionally phase-bright and lysed (**Fig. 7**). This series of experiments in the Δ*cpsE* background suggests that CpsA can partially substitute for LytR and Psr functions.

**Fig. 7.**
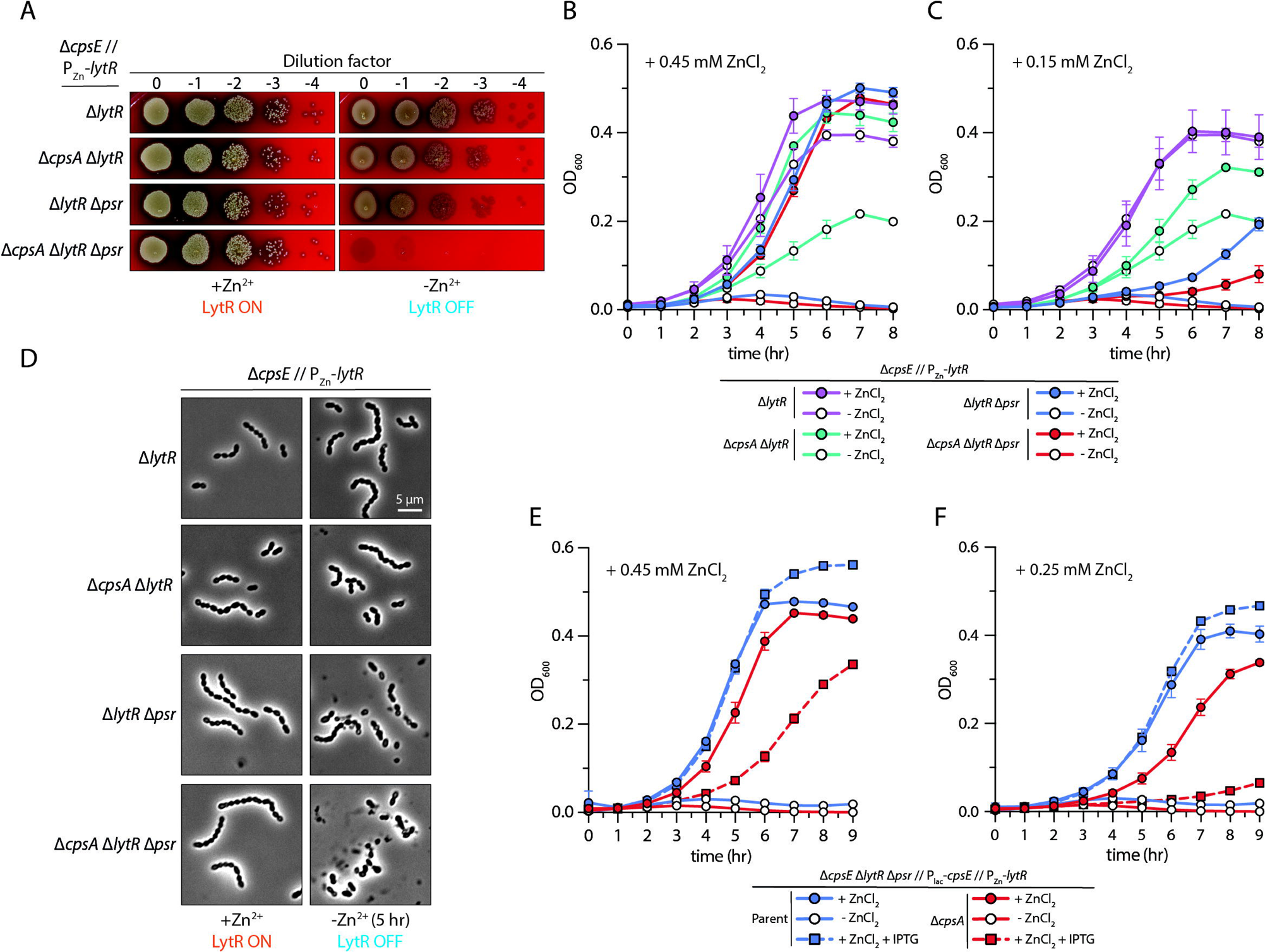
LCP ligases are semi-redundant in *S. pneumoniae* strain D39W. (**A**) *cpsA* promotes growth in strains lacking Δ*lytR* and Δ*psr*, independent of capsule production. Strain NUS5935 [Δ*cpsE* // P_Zn_-*lytR*] and its derivatives lacking the indicated genes were grown in BHI supplemented with 0.45 mM Zn^2+^ at 37 °C in 5% CO_2_. Cultures were normalized to the same optical density, serially diluted, and spotted onto blood agar plates with or without 0.45 mM Zn^2+^. Plates were imaged after overnight incubation at 37 °C in 5% CO_2_. Shown are representative images from three biological repeats. The strains described in (**A**) were grown in BHI in the presence of 0.45 mM Zn^2+^ until the cultures reached exponential phase, washed twice in BHI, then normalized to the same optical density in BHI with or without 0.45 mM ZnCl_2_ (**B**) or 0.15 mM ZnCl_2_ (**C**) and let grow at 37 °C in 5% CO_2_. Growth was monitored by measuring the optical densities of the cultures over time. Plotted are the means and standard deviations from three biological repeats. (**D**) Five hours after subculturing, cells were observed under phase-contrast microscopy. Bar, 5 µm. Shown are the representative micrographs from three biological repeats. (**E**) When CpsA is inactivated, capsule precursors compete with wall teichoic acid precursors for LytR. Strains NUS7125 [Δ*cpsE* Δ*lytR* Δ*psr* // P_lac_-*cpsE* // P_Zn_-*lytR*] and NUS7126 [Δ*cpsA* Δ*cpsE* Δ*lytR* Δ*psr* // P_lac_-*cpsE* // P_Zn_-*lytR*] were grown in BHI supplemented with 0.45 mM Zn^2+^ at 37 °C in 5% CO_2_. Cultures were washed twice, normalized to the same optical density, and inoculated in BHI with or without 0.45 mM ZnCl_2_ (**E**) or 0.25 mM ZnCl_2_ (**F**). IPTG was added where indicated to induce capsule production. Growth was monitored by measuring the optical densities of the culture over time. Plotted are the means and standard deviations from three biological replicates.

We then examined the effect of capsule expression in strains with reduced LCP activity. These experiments were done in the NUS7030 [Δ*cpsE* // P_lac_-*cpsE*] background. This allowed us to control capsule expression with IPTG while maintaining cell viability, where *lytR* is under the control of a zinc-inducible promoter at an ectopic locus [Δ*cpsE* Δ*lytR* // P_lac_-*cpsE* // P_Zn_-*lytR*]. We also inactivated *psr* in the resulting strain. Under this condition, cells were unable to grow in BHI when *lytR* was depleted (**Fig. 7**). Inducing capsule production did not exacerbate the phenotype, unless *cpsA* was simultaneously inactivated (**Fig. 7**). This effect was more pronounced when the *lytR* expression was reduced by decreasing the concentration of the Zn^2+^ inducer in the medium (**Fig. 7**). Our data indicate that when LytR is the only LCP ligase in the cell, capsule production becomes lethal, possibly because the capsule polymers compete with the WTA. The latter is thought to be required for growth (75).

The lethality of Δ*walK* Δ*cpsA* allowed us to identify essential residues of CpsA through mutagenesis sequencing (Mut-seq) (18, 76). First, we PCR-mutagenized *cpsA* and introduced it via allelic exchange into a strain in which the *cpsA* gene was replaced with the sweet Janus cassette (P-*sacB*-*kan*-*rpsL*^+^) (77). If the mutagenized *cpsA* was incorporated, the strain would become resistant to sucrose and streptomycin, generating a library of strains carrying different CpsA variants. Yet, despite numerous attempts, we could not build a library of meaningful size in the NUS6142 background [*rpsL1* Δ*cpsA*::P-*sacB*-*kan*-*rpsL*^+^ Δ*cpsE* Δ*walK* // P_Zn_-*cpsE*], likely because Δ*walK* reduced natural competence (78). To overcome the low transformation efficiency of the Δ*walK* mutant, we made the library in the isogenic *walK*^+^ strain (NUS0546 [*rpsL1* Δ*cpsA*::P-*sacB*-*kan*-*rpsL*^+^ Δ*cpsE* // P_Zn_-*cpsE*]), generating approximately 6 million transformants. Next, we pooled these mutants and transformed ≈4 million cells with a genetic cassette to inactivate *walK*, resulting in ≈22 million transformants. Colonies were pooled, serially diluted, and plated on blood agar plates in the presence or absence of Zn^2+^ inducer. Unless the cell harbors a functional copy of *cpsA*, induction of *cpsE* expression is lethal. Consistently, we observed nearly a two-fold reduction in plating efficiency when Zn^2+^ was added at a final concentration of 0.45 mM, and an additional two-fold decrease when the concentration increased to 0.85 mM. Survivors on plates with or without Zn^2+^ were collected, and the *cpsA* alleles were sequenced. We found several residues in CpsA that appeared immutable in the non-permissive condition (**Fig. S7, table S4**). Among them, residues R244, D268, I273, L339, and E363 appeared to be indispensable for CpsA function, as changing them to alanine disrupted CpsA function (**Fig. S7**). Residue D268 is one of the potential catalytic bases of CpsA, although it is ∼5 Å away from the predicted position of the incoming nucleophile. Moreover, R244 is part of the arginine pairs that coordinate the pyrophosphate head group of C55-P. (28) Thus, the five residues we identified as essential in CpsA are likely clustered around the active site of the conserved extracellular LCP domain.

Next, we tested whether CpsA alone is sufficient to support growth in the absence of other LCP ligases. To do this, we constructed a strain lacking *cpsE* and the three LCP genes *cpsA*, *lytR*, and *psr* (Δ*lcp*). Cell viability was maintained by expressing *lytR* in trans [P_Zn_-*lytR*]. Under these conditions, cells could not grow unless the inducer was added to the medium (**Fig. 8**). As a positive control, the introduction of a constitutively expressed copy of *lytR* (P*-lytR*) restored growth. Similarly, constitutive expression of *cpsA*, but not as much with a *cpsA* hypomorph (*cpsA*^R244A^), also alleviated the growth defects in the Δ*lcp* strain (**Fig. 8**). To rule out that the restored growth was caused by leaky *lytR* expression, we deleted the P_Zn_-*lytR* cassette, resulting in a strain in which CpsA is the only LCP enzyme. Deleting the same cassette in cells producing the CpsA^R244A^ variant generated no viable colonies, indicating CpsA activity was required for growth. Although the resulting strain survived, the cells formed chains (**Fig. 8** and **S8**), suggesting partial complementation. We conclude that CpsA can support growth as the sole LCP enzyme when mildly overexpressed. It is likely because CpsA compensates for the functions of LytR and Psr, thereby restoring WTA attachment to PG.

**Fig. 8.**
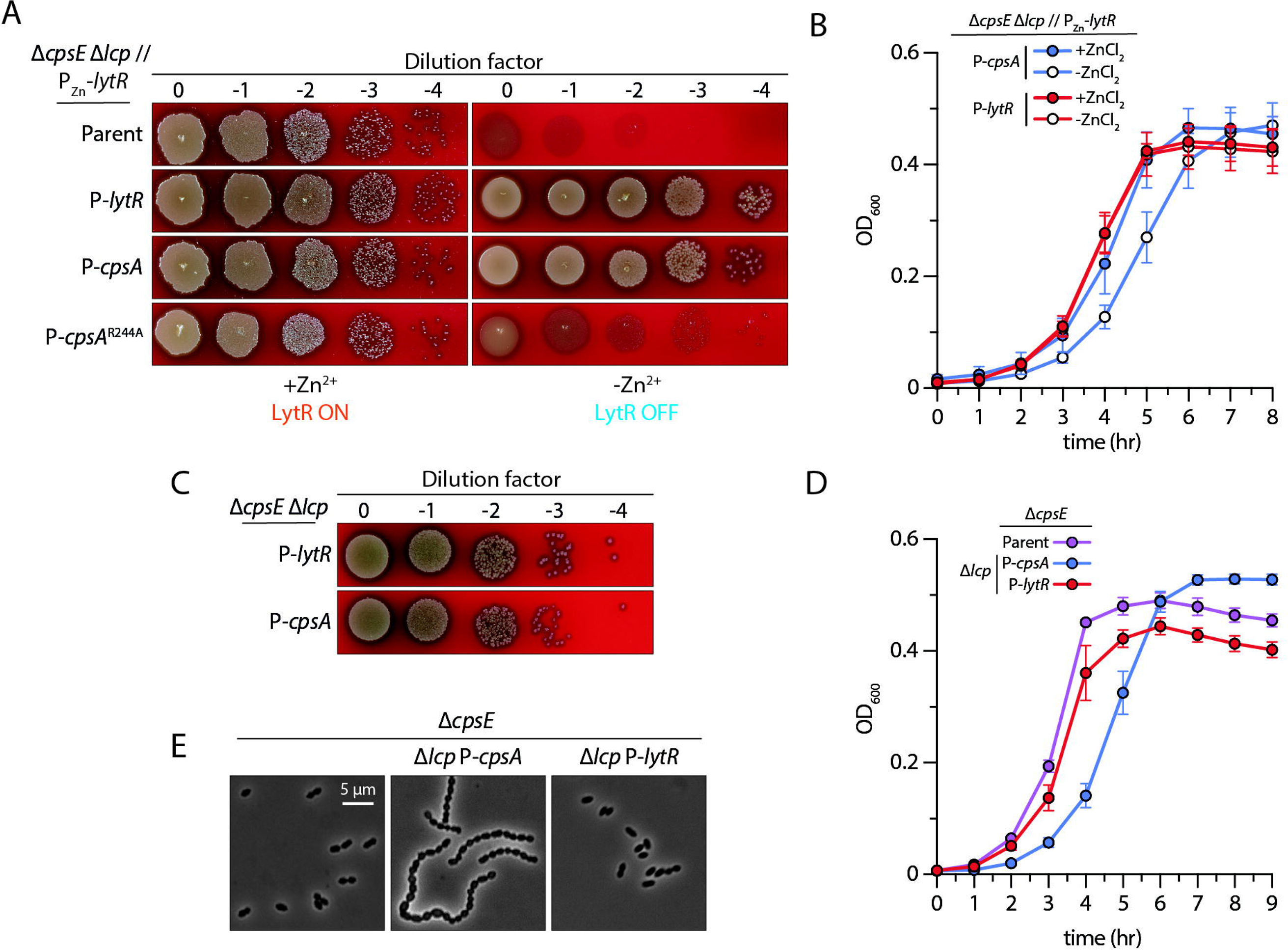
Semi-redundancy of LCP ligases in *S. pneumoniae*. (**A**) Overproducing CpsA bypasses the requirement of other LCP ligases for growth. Strain NUS6195 [Δ*cpsE* Δ*lcp* // P_Zn_-*lytR*] and its derivatives expressing the indicated proteins (LytR, CpsA, or CpsA^R244A^) were grown in BHI supplemented with 0.45 mM Zn^2+^ at 37 °C in 5% CO_2_. Cells were washed twice with BHI without added Zn^2+^ and then resuspended in BHI. After the suspensions were normalized by their optical density, they were serially diluted and spotted onto blood agar plates with or without supplemented Zn^2+^. Plates were visualized after incubating overnight at 37 °C in 5% CO_2_. (**B**) Cells depleted of all LCP ligases, except CpsA, are impaired in growth. Strains NUS6907 [Δ*cpsE* Δ*lcp* // P_Zn_-*lytR* // P-*cpsA*] and NUS6908 [Δ*cpsE* Δ*lcp* // P_Zn_-*lytR* // P-*lytR*] were grown in BHI supplemented with Zn^2+^ at 37 °C in 5% CO_2_. Cells were washed twice with BHI, and the cultures were normalized by their optical densities. Growth was monitored by measuring OD_600_ over time. Plotted are the means and standard deviations from three biological replicates. (**C**) CpsA can substitute for the function of LytR and Psr. Strains NUS7288 [Δ*cpsE* Δ*lcp* // P-*cpsA*] and NUS7289 [Δ*cpsE* Δ*lcp* // P-*lytR*] were grown, spotted on blood agar plates, and imaged as described in (**B**). (**D**) Strains indicated in (**C**) were grown in BHI to the early exponential phase and imaged by phase-contrast microscopy (**E**). Scale, 5µm. Experiments were performed three times with similar results.

## DISCUSSION

The capsule polysaccharide plays a crucial role in pneumococcal virulence. The first four genes in the capsule locus, *cpsABCD*, are highly conserved across serotypes. Their exact functions in capsule synthesis remain incompletely understood. Our previous studies have demonstrated that the CpsBCD bacterial tyrosine kinase system regulates polysaccharide chain length and the site of capsule synthesis (22). CpsCD is directed to the septum by the divisome, which then recruits other capsule enzymes to the mid-cell (24). In this study, we continued to investigate the functions of *cpsABCD* by identifying genetic interactions involving these conserved genes. We revealed that one of the essential roles of CpsD is to facilitate CpsC-CpsH interaction. In addition, *walK* is essential for growth in *cpsA* and *lytR* cells. We also experimentally confirmed the functional redundancy of LCP ligases. Additionally, cells lacking PG-modifying enzymes such as OatA and PgdA are resistant to the defects caused by Δ*cpsA* Δ*walK*. These findings support a working model in which WTA and other PG modifications influence PG hydrolase activities. These changes are likely sensed by the WalRK two-component system, which maintains the intricate balance of PG synthesis and remodelling by positively regulating PG hydrolases. Otherwise, cells cannot survive unless PG hydrolytic activity is restored by inactivating PG-modifying enzymes OatA and PgdA.

### CpsD primarily functions as a scaffolding protein bridging CpsC and CpsH

Coordinating the synthesis of different layers of the cell envelope is required for survival. Here, we demonstrate that the essentiality of CpsD for growth can be bypassed by bridging the interaction between CpsH and CpsC. This interaction appears crucial for maintaining sufficient capsule polymerase activity. Without it, C55-P recycling may be halted due to the accumulation of unpolymerized capsule repeating units. As expected, *cpsC* and *cpsD* have negative genetic interactions with genes involved in C55-P synthesis, such as *ispA* and *panT*. In contrast, we did not observe significant genetic interactions involving *cpsB*. Phosphorylation of CpsD is thought to reduce CpsC-CpsH interaction (22). Thus, Δ*cpsB* likely boosts CpsH activity, which aligns with the shorter capsule polymers seen in cells lacking CpsB phosphatase activity. The signal detected by the CpsBCD system that regulates CpsD-P levels remains unknown. We speculate that it may be oxygen (79–82), as its levels vary widely across body parts. Oxygen levels may therefore serve as a marker of different host niches, allowing the cell to adjust the length of its capsule chain accordingly. Consistently, pneumococcal cells produce a thicker capsule in blood than in growth media or under conditions that simulate the respiratory tract (83).

### The synthetic lethal relationship between *cpsA* and *walK*

We found that *cpsA* does not share the same set of genetic interaction partners as *cpsC* and *cpsD*. Instead, it forms a synthetic lethal pair with *walK*. The WalRK TCS may sense cell envelope stress induced by Δ*cpsA*, likely due to reduced PcsB activity resulting from reduced WTA levels or accumulation of Und-PP-linked WTA precursors. In response, WalRK adjusts the levels of *pcsB* and other PG hydrolases to correct these deficiencies. We speculate that a possible signal of WalK is the presence of WTA intermediates on the cytoplasmic membrane, because WalK in pneumococcus lacks the sCache domain required to detect PG hydrolytic products. However, since loss of WTA from the cell surface likely has pleiotropic effects across the cell envelope, we cannot rule out that any of these feed back to regulate WalK. We did not detect synthetic lethality or sickness when we combined Δ*walK* with Δ*cpsC* or Δ*cpsD*, which argues against direct WalRK sensing of Und-P levels (22).

The WalRK system is required for viability when LCP ligase activity is reduced (**Fig. 4** and **5**). The requirement for PcsB upregulation to suppress Δ*cpsA* Δ*walK* lethality is consistent with a recent report that in *Bacillus subtilis*, deletions of LCP ligases decrease cell wall hydrolase activity (74). Since WTA are thought to direct proper peptidoglycan remodeling by localizing cell wall hydrolases (72), unligated lipid-linked precursors on the outer leaflet of the cytoplasmic membrane may lead to a deficiency in peptidoglycan hydrolysis, rendering the PG hydrolase LytE required for growth (74). In the pneumococcal system, WalK might sense lipid-linked intermediates at the inner leaflet via its cytoplasmic domain and restore the balance between secondary polymer attachment and peptidoglycan remodeling, although we have not ruled out the possibility that the WalRK system senses signals indirectly via StkP (84) or other factors. In addition to the pneumococcal WalK, loss of LCP ligases in *S. aureus* (MsrR, SA0808, and SA2103) (85) and *B. subtilis* (ΔTagTV) (74) also triggers cell wall stress responses via WalRK signaling. In the latter scenario, stress is alleviated by inactivating *tkmA*, a PCP that facilitates the synthesis of an unknown polymer (74).

### LCP enzymes are semi-redundant and interchangeable

The LytR/Cps2A/Psr (LCP) ligases are found in virtually all Gram-positive bacteria (86). Phylogenetic analysis reveals their ancient origin and subsequent diversification (32). The conservation of catalytic residues across species, combined with variable loop regions, suggests that LCP proteins evolved to accommodate different secondary wall polymer donors while maintaining a common catalytic mechanism (32). Our work provides genetic evidence supporting CpsA can ligate CPS and WTA to PG, circumventing the cell lysis issue previously reported in the Δ*cpsA* Δ*lytR* mutant (41). The substrate preference of CpsA can be overcome by its overexpression, thereby bypassing the requirement of LytR and Psr for growth (**Fig. 8**). As previously reported (39, 40), capsule ligation was not disrupted in strains lacking CpsA, presumably because the capsule precursors are conjugated to PG by LytR and Psr. Thus, in the presence of WalK, the three LCP paralogs in pneumococcus can ensure the attachment of cell wall polymers, even when one of them is inactivated (41).

LytR and Psr are thought to be WTA ligases that attach teichoic acid polymers to the C6 position of the N-acetyl muramic acid (MurNAc) residue (34, 75). If they also attach capsule polymers, we expect them to do so at the same sugar residue, thereby changing the CPS attachment site (21). Similarly, if CpsA attaches WTA polymers to PG, it should link them to the GlcNAc residue rather than to MurNAc. If our hypothesis is true, it suggests that CpsA has a different preference for attachment sites than LytR and Psr, possibly to prevent competition for the acceptor between CPS and WTA polymers. This is intriguing because purified LCP enzymes can attach secondary wall polymers to chitin (33), indicating they lack strict acceptor specificity. The strain expressing CpsA as its only LCP ligase still formed chained cells in liquid culture (**Figs. 8** and **S8**). This result has at least two implications. CpsA might prefer CPS over WTA polymers. Although the P-*cpsA* cassette probably increased CpsA expression, it remains insufficient to overcome the specificity barrier. Another possibility is that the WTA attachment site may be important for its function. Attaching it to GlcNAc or changing the presumed linkage type might partially reduce its function.

The interchangeability of LCP enzymes suggests that they also lack strong donor specificity. Crystal structures of Psr (87) and other LCP proteins (88) indicate that the loops of the polysaccharide binding domain are flexible, which may contribute to the broad range of substrates that can be accommodated into the active site (28, 30, 31, 89, 90). However, when all LCP ligases are present, substrate preference appears to emerge (**Fig. 6** and **7**). This substrate preference may also stem from the subcellular localization of the ligases and enzymes that generate the corresponding polymers (24, 25). Additionally, CpsA is thought to form a direct glycosidic linkage rather than a phosphodiester bond (21). If it can substitute for the functions of LytR and Psr, it should also link WTA to PG via a direct glycosidic bond, suggesting that the phosphodiester bond is not a strict requirement for WTA function. Similarly, if LytR and Psr attach CPS to PG, we speculate that the attachment is via a phosphodiester linkage. How this change affects CPS functions remains to be elucidated. An alternative hypothesis is that CpsA in *S. pneumoniae*, like all other LCP ligases, catalyses the formation of a phosphodiester linkage (26). It is possible because a close homolog of CpsA in *Streptococcus agalactiae* serotype 3 (GBSIII) is known to make a phosphodiester bond between the capsule polymer and the GlcNAc residue of PG. These two proteins are ∼50% identical and ∼70% similar, with greater homology in the C-terminal LCP domain. Future studies are required to examine these mutually exclusive possibilities, which will likely require sophisticated mass spectrometry experiments. Moreover, it remains to be determined whether changes in polymer attachment will have functional consequences in laboratory conditions or in the host.

### Connections Between PG Modifications and LCP Function

The suppression of Δ*cpsA* Δ*walK* lethality by Δ*pgdA* and Δ*oatA* deletions indicates connections between peptidoglycan modifications and LCP function. PG O-acetylation by OatA and N-deacetylation by PgdA have been shown to increase lysozyme resistance (91). This notion is supported by multiple reports indicating that PG modifications reduce PG hydrolysis. For example, the PG deacetylase PdaC in *B. subtilis* is thought to reduce the activities of LytE (92) and CwlO (66). WalRK also regulates PdaC levels, providing an additional layer of control for the system (66). Additionally, O-acetylation also affects the accessibility of attachment sites for LCP substrates at the C6 position of MurNAc. Therefore, inactivating *oatA* may limit the amount of lipid-linked WTA precursors on the cell surface. Indeed, Δ*oatA*, but not Δ*pgdA*, restored the chaining phenotype of the Δ*walK* mutant (**Fig. 6**). Consistent with the chaining phenotype related to PG hydrolysis and WTA attachment, this defect could also be corrected by ectopically expressing PcsB or LytR (**Fig. 6**). Moreover, depletion of *pcsB* also results in the formation of chains of cells, likely due to the inability to split the septum after cell division (73). Thus, in addition to controlling the WTA amount on the cell surface through cleavage (72), OatA may block WTA attachment by installing an acetyl group on PG.

Based on these findings, we propose that CpsA can function as a capsule ligase and that LytR and Psr can compensate for its absence (**Fig. 9**). However, involving LytR and Psr in ligating CPS polymers will inevitably divert resources from the WTA pathway, leading to cell envelope stress, including the accumulation of lipid-linked WTA precursors. The WalRK system detects this stress and responds by increasing the production of PG hydrolases. In this scenario, sensing is likely mediated by the cytoplasmic PAS domain of the WalK histidine kinase, since its extracellular domain contains only 12 amino acids (55). Furthermore, WalRK in *S. pneumoniae* lacks the auxiliary regulatory factor WalHI (55), suggesting that signal sensing is likely performed directly by WalK (55). The synthetic lethality between LCP ligases and WalK suggests that inhibiting both polymer attachment and cell envelope stress responses could provide synergistic therapeutic effects. Additionally, the mechanism by which WalRK detects WTA deficiency remains unclear and will be a subject of future studies.

**Fig. 9.**
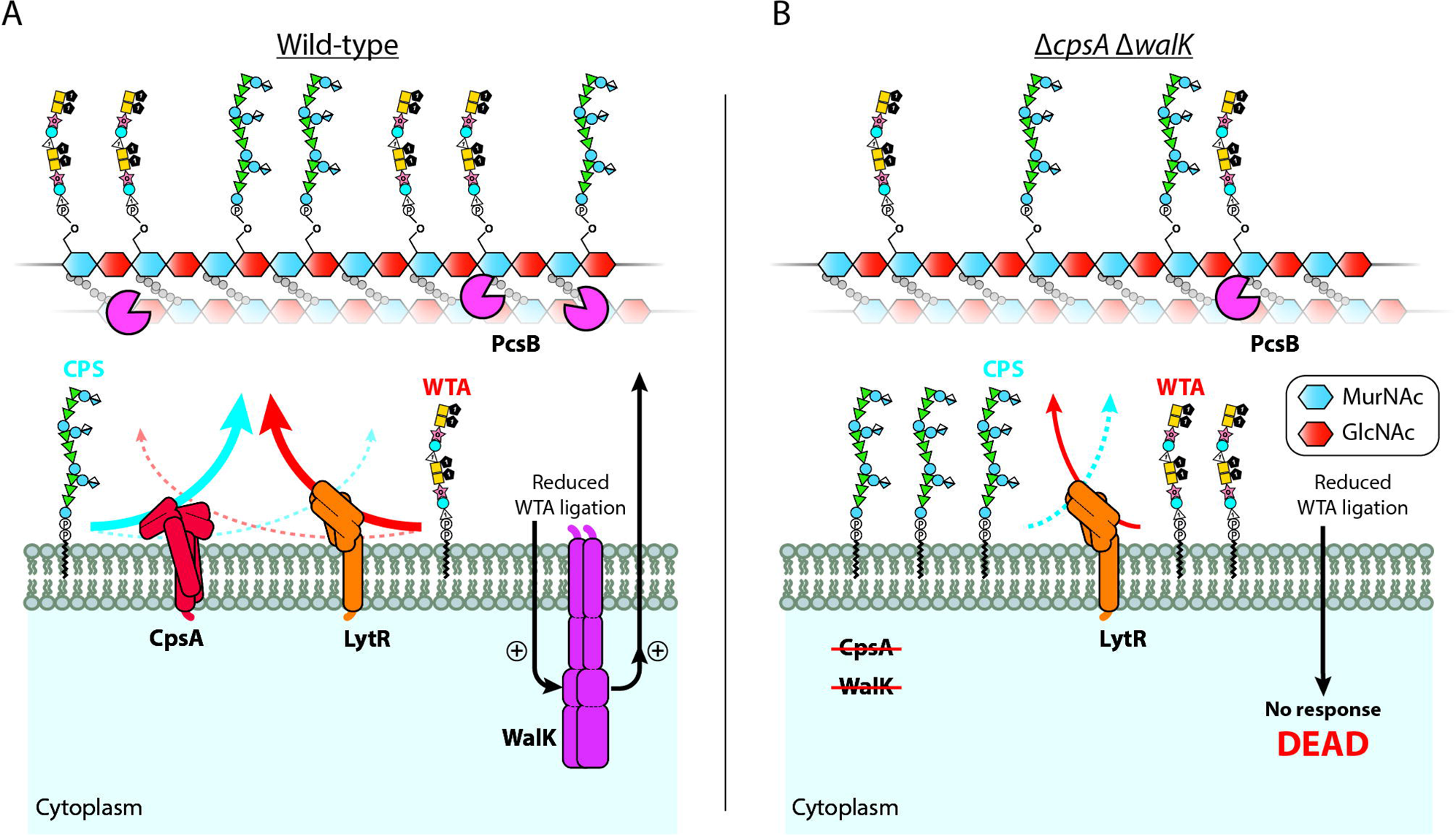
The WalRK system responds to defects caused by reduced LCP ligase activity. Shown is the working model of a compensatory mechanism that maintains capsule synthesis when the candidate capsule ligase CpsA is absent. (**A**) In wild-type cells, CpsA attaches capsule polymers to the GlcNAc residues of PG, while LytR attaches WTA polymers to the MurNAc residues. For simplicity, we assume that the linkage between PG and secondary polymers is a phosphodiester bond. This hypothesis needs further experimental validation. When WTA attachment is insufficient, WalK detects this stress and responds by changing gene expression, such as increasing PcsB levels. (**B**) When *walK* and *cpsA* are inactivated, LytR attaches both capsule and WTA polymers to PG. As a result, the cell accumulates lipid-linked precursors and has reduced WTA on its surface. This causes cell envelope stress that cannot be alleviated by the WalRK two-component system. Consequently, the cell cannot survive due to a lack of PG hydrolytic activity.

## MATERIALS AND METHODS

### Bacterial strains and growth conditions

The strains used in this study are listed in **Table S1**. Unless otherwise specified, *S. pneumoniae* cells were grown in brain heart infusion broth (BHI) or on tryptic soy agar plates supplemented with 5% (v/v) sheep blood (blood plates) (Biomed Diagnostics, 221261) at 37°C in 5% CO_2_. Where indicated, antibiotics were added at final concentrations of 0.3 μg/mL for erythromycin (Erm), 300 μg/mL for kanamycin (Kan), 300 μg/mL for streptomycin (Str), 200 μg/mL for gentamycin (Gent), 2.5 µg/mL for chloramphenicol (Chlor), and 200 μg/mL for spectinomycin (Spec). Expression of the *czcD* promoter (P_Zn_) was induced by adding ZnCl_2_ to a final concentration between 150 μM and 850 µM as specified for each figure. To counteract the cell division defects caused by Zn^2+^ toxicity, 1:10 of the amount of MnCl_2_ was added to the medium simultaneously (93, 94). Expression of the IPTG-inducible promoter (P_lac_) was induced by adding IPTG to a final concentration of 1 mM in cells constitutively expressing the *lacI* repressor gene (95).

### Strain construction and transformation

Primers for generating PCR amplicons are listed in **Table S2**. PCR products were synthesized using Phusion DNA polymerase. Genetic cassettes were assembled using Gibson assembly (96). Pneumococcal strains were constructed by transforming cells with PCR amplicons, assembled fragments, or genomic DNA (gDNA) after inducing natural competence, as described previously (97, 98). Briefly, 1 mL of log phase culture (OD_600_ = 0.1 to 0.4) was incubated in BHI containing 1 mM CaCl_2_, 0.04% (w/v) bovine serum albumin, and 250 ng of competence-stimulating peptide (CSP-1) for 10 minutes at 37°C in 5% CO_2_. DNA was added, and the cells were incubated for 1.5 hours at 37°C in 5% CO_2_ before being plated on blood agar supplemented by the indicated antibiotics. Plates were incubated overnight at 37°C in 5% CO_2_. Markerless deletion mutants were generated using the Janus (P-*kan*-*rpsL*^+^) or Sweet Janus (P-*sacB*-*kan*-*rpsL*^+^) cassettes, or derivations thereof, using alternative antibiotic markers, with counter-selections using Str and 5% (w/v) sucrose (77, 99). gDNA was extracted using the DNeasy Blood & Tissue kit (Qiagen) as per the manufacturer’s instructions.

### Measurements of growth

For spot dilutions, strains were grown in the indicated media at 37°C in 5% CO_2_ until the OD_600_ was between 0.1 and 0.5. Unless otherwise specified, cultures were diluted to an OD_600_ of 0.05 and ten-fold serially diluted. Two microliters of each dilution were spotted onto blood agar plates with or without added ZnCl_2_ and MnCl_2_. Plates were incubated overnight before imaging. For growth curves, strains were grown in the indicated media at 37°C in 5% CO_2_ until the OD_600_ was between 0.1 and 0.5. Cultures were diluted to an OD_600_ of 0.01 with BHI broth in a final volume of 200 µL before being transferred to 96-well plates. If the cells were initially grown in broth supplemented with ZnCl_2_ and MnCl_2_, they were washed twice with 1 mL of plain BHI and diluted to an OD_600_ of 0.01 in the indicated growth medium to a final volume of 200 µL. Growth measurements were performed by either (i) incubating the plate in a Tecan Spark 10M microplate reader at 37°C and taking OD_600_ measurements every 20 minutes after shaking for 20 seconds or by (ii) incubating at 37°C in 5% CO_2_ in the incubator and transferring the plate to a Tecan Spark 10M microplate reader, shaking for 20 seconds, and measuring the OD_600_ over time.

### RB Tn-seq and BarSeq analysis

The indicated strains were grown in plain BHI at 37°C in 5% CO_2_ until the cultures reached the log phase. Cells were normalized to an OD_600_ of 0.05 in plain BHI before being transformed with 300 ng of RBloxSpec transposon library gDNA per mL of culture after inducing natural competence (see Strain construction and transformation) (47). Transformed cells were plated onto blood plates supplemented with 450 µM ZnCl_2_, 45 µM MnCl_2_, and spectinomycin and incubated overnight at 37°C in 5% CO_2_. Over 500,000 transformants were obtained per strain. To collect the cells, plates were flooded with 3 mL of BHI, scraped with a spreader, and the cell suspensions were poured into a collection tube. BarSeq to amplify the barcodes was performed on 200-300 ng of genomic DNA prepped from the collected cells as described previously (47), using primers O1373 and one of the indexed reverse primers. Insertions were quantified and used to calculate gene fitness (the estimated log2 changes in the abundance of mutants of each gene) and significance (t) scores using Perl scripts described previously (46). Here, fitness was calculated by comparing the recovered Tn insertions between the parent strain (NUS0267 [Δ*cpsE* // Δ*bgaA*::P_Zn_*-cpsE*]) and its derivatives (e.g., NUS0796 [Δ*cpsA* Δ*cpsE* // Δ*bgaA*::P_Zn_*-cpsE*) after transforming them with the genomic DNA purified from the Tn library and selecting the mutants on plates supplemented with ZnCl_2_ and MnCl_2_ (see above). For isolating suppressors of the Δ*cpsA* Δ*walK* interaction, NUS3698 [Δ*cpsE* Δ*cpsA* Δ*walK*::P*-erm-rpsL* // Δ*bgaA*::P_Zn_*-cpsE*) was transformed with 400 ng of genomic DNA purified from the RBloxSpec transposon library, plated on blood plates without added ZnCl_2_ and MnCl_2_, and incubated overnight at 37°C in 5% CO_2_. This library was collected, plated onto blood plates with or without 450 µM ZnCl_2_ and 45 µM MnCl_2_, incubated overnight at 37°C in 5% CO_2,_ and collected and quantified as above, using primers O1144 and one of the indexed reverse primers (**Table S2**) for sequencing. The fitness and t values are listed in **Table S3**.

### Quantitative reverse transcription PCR (qRT-PCR)

To quantify the relative expression of *pcsB*, the indicated strains were grown to the log phase in BHI at 37°C in 5% CO_2,_ then normalized to an OD_600_ of 0.1 in the same medium. Cells were then induced with 450 µM ZnCl_2_ and 45 µM MnCl_2_ and incubated for one hour at 37°C in 5% CO_2_. Before and after induction, 500 µL of culture was removed and mixed with 1 mL of RNAprotect Bacteria Reagent (Qiagen), vortexed, and incubated at room temperature for 5 minutes. Samples were centrifuged for 10 minutes at 5,000 x *g*, the supernatant was removed, and pellets were stored at −80°C. RNA was extracted using the RNeasy Mini Kit (Qiagen) according to the manufacturer’s instructions for enzymatic lysis and proteinase K digestion of bacteria (Protocol 4), with added supplementation of 40 U of mutanolysin at the lysis step. Purified RNA was then treated with DNase I using the DNA-free™ DNA Removal Kit (Invitrogen) according to the manufacturer’s instructions. Treated RNA samples were normalized to 15 ng/µL and converted to cDNA using the High-Capacity cDNA Reverse Transcription Kit (Thermo Fisher) according to the manufacturer’s instructions. qPCR was performed on 1:20 dilutions of cDNA samples using the FastStart Essential DNA Green Master kit (Roche), using primers O5195/O5196 for *pcsB* and O5199/O5200 for *rpoB* (control). Reactions were run on a LightCycler 96 (Roche) with the following settings: 95°C for 10 minutes; 45 cycles of 95°C for 10 seconds, 55°C for 10 seconds, 72°C for 5 seconds; and a final melting step of 95°C for 10 seconds, 65°C for 60 seconds, 97°C for 1 second. Relative abundance was determined by the ΔΔCt method (100) and normalized to the copy number of *rpoB*.

### Flow cytometry

Strains were grown in BHI at 37°C in 5% CO_2_ until the OD_600_ was between 0.05 and 0.2. Cells were subsequently inactivated by incubation in 1% paraformaldehyde for one hour. For induced samples, cells were grown in the indicated concentration of ZnCl_2_/MnCl_2_ directly or first grown in plain BHI to log phase, normalized to an OD_600_ of 0.01 in BHI with ZnCl_2_/MnCl_2_, and then grown for five doublings (to OD_600_ = 0.3) at 37°C in 5% CO_2_ before paraformaldehyde inactivation. Inactivated cells were directly diluted into 1x phosphate-buffered saline (PBS) and analyzed using a Becton-Dickinson Fortessa at the Flow Cytometry Laboratory in the Life Sciences Institute Immunology Programme at the National University of Singapore. The forward scatter (FSC) and side scatter (SSC) were measured for 10,000 events per sample. Floreada.io (https://floreada.io) was used to gate the samples, while FlowJo™ (BD) was used to generate and export the figures.

### Capsule immunoblotting

The indicated strains were grown in BHI at 37°C in 5% CO_2_ until the OD_600_ was between 0.1 and 0.5. Cells were normalized to an OD_600_ of 0.1, supplemented with 450 µM ZnCl_2_ and 45 µM MnCl_2_, and grown for 1.5 hours at 37°C in 5% CO_2_. Cells were normalized by OD_600,_ centrifuged at 16,000 x *g* for 10 minutes, and the supernatant was removed. Cells were resuspended in 100 µL 1x Laemmli buffer with 2 µL proteinase K (Qiagen, 19133) and incubated for 1 hour at 56°C. Fractionation of protoplasts and cell wall fragments was performed similarly to previously (72) with slight modification. Briefly, following the induction of CPS production as above, cultures were normalized to the same OD_600,_ centrifuged at 5,000 x *g* for 5 minutes, and resuspended in 1/10 of the volume of SMM buffer (0.5 M sucrose, 20 mM maleic acid, 20 mM MgCl_2_, pH 6.5). 75 U/mL of mutanolysin (Sigma, M9901), 5 mg/mL lysozyme (GoldBio, L-040), and 12.5 µg/mL of purified LytA (50) were added to each sample. Samples were incubated overnight on a rotator at low speed at room temperature. Protoplast formation was confirmed via phase-contrast microscopy. A “combined” fraction was collected, and the remaining samples were centrifuged at 5,000 x *g* for 5 minutes to pellet the protoplasts. The supernatant was removed, filtered through a 0.22 µm filter, and collected (“cell wall” fraction). The remaining pellet was washed once with an equal volume of SMM buffer, resuspended in the same volume of SMM buffer, and collected (“protoplast” fraction). Samples were mixed with an equal volume of 1x Laemmli buffer with 2 µL of proteinase K and incubated for 1 hour at 56°C. All samples were separated on 10% SDS-PAGE gels and transferred to polyvinylidene fluoride (PVDF) membranes. Membranes were blocked for 30 minutes at room temperature in blocking buffer (1x PBS with 0.05% [v/v] Tween 20 and 5% [w/v] skimmed milk). Blots were probed with anti-serotype 2 CPS antiserum (SSI Diagnostica) at a 1:5,000 dilution overnight at 4°C in blocking buffer, washed twice with PBST (1x PBS with 0.05% [w/v] Tween 20), and probed with goat anti-rabbit horseradish peroxidase secondary antibodies (Thermo Fisher, A16110) in blocking buffer at a 1:10,000 dilution for one hour at room temperature. The membranes were washed twice with PBST and developed using the SuperSignal™ West Pico PLUS Chemiluminescent Substrate (Thermo Fisher).

### Live/dead staining and microscopy

The indicated strains were grown in plain BHI at 37°C in 5% CO_2_ until the OD_600_ was between 0.1 and 0.5. Cells were normalized to an OD_600_ of 0.01, supplemented with or without the stated concentration of ZnCl_2_/MnCl_2_, and grown for five doublings (to OD_600_ = 0.3) at 37°C in 5% CO_2_ before visualization. Cells were collected by centrifugation at 8,000 x *g* for 2 minutes, washed twice in 150 mM NaCl, and resuspended in the same solution. Cells were stained using the LIVE/DEAD BacLight Bacterial Viability Kit (L7007; Molecular Probes, Invitrogen) according to the manufacturer’s instructions, and placed on 1% (w/v) agarose pads for visualization. For all microscopy images, cells were imaged using a Nikon ECLIPSE Ti2 inverted microscope equipped with an Andor Sona-4BV11 sCMOS camera (Oxford Instruments) and an X-Cite XYLIS XT720S LED light source (Excelitas).

### Mutagenesis sequencing (Mut-seq)

The native locus of *cpsA* was PCR-mutagenized by amplifying with primers N1094 and N1095 using GoTaq DNA polymerase (M7123; Promega) and the following thermocycler settings: 95°C for 4 minutes; 20 cycles of 95°C for 30 seconds, 52°C for 30 seconds, 72°C for 1.5 minutes; and finally 72°C for 7 minutes. PCR products were purified and concentrated using the Qiagen PCR purification kit. NUS0546 [Δ*cpsE* Δ*cpsA*::P*-sacB-kan-rpsL* // Δ*bgaA*::P_Zn_*-cpsE*] was transformed with the PCR mutagenized DNA fragment. Transformants were selected on plates supplemented with streptomycin and sucrose as described above (see Strain construction and transformation). The library contains ∼6.5 million transformants. Cells were pooled and stored at −80°C in 15% (v/v) glycerol. This library was thawed and transformed with Δ*walK*::P*-erm-rpsL,* amplified from strain NUS3639 with primers O3587 and O3590 using Phusion polymerase, yielding 22 million transformants. Cells were collected, pooled, and stored at −80°C in 15% (v/v) glycerol. This resultant library was thawed and plated on blood agar with or without 850 µM ZnCl_2_ and 85 µM MnCl_2,_ and incubated overnight at 37°C in 5% CO_2_, yielding 16 million colony-forming units (CFUs) for the ZnCl_2_-treated condition and 53 million CFUs for the uninduced condition. Cells were collected, and the DNA was extracted. The *cpsA* locus was amplified with primers N1094 and N1095 using Phusion polymerase in 9 total PCR reactions with 300 ng DNA per reaction using the following thermocycler settings: 95°C for 2 minutes; 25 cycles of 95°C for 30 seconds, 50°C for 30 seconds, 72°C for 1 minute; and finally 72°C for 7 minutes. Amplicons were fragmented and sequenced at Azenta Life Sciences using the NovaSeq 150PE platform. Data analysis was performed using the CLC Workbench software (Qiagen) as described previously (18). The raw data of the Mut-seq experiment are listed in **Table S4**.

## ACKNOWLEDGEMENTS

We thank members of the L.-T.S. laboratory for their helpful discussions. The P-*lacI*(rev)-P_lac_ cassette was generously provided by the laboratory of Dr. Jan-Willem Veening. We also thank the staff at the Flow Cytometry Laboratory in the Life Sciences Institute Immunology Programme at the National University of Singapore for support.

## Funding

This work was supported by grants from:

The National Research Medical Council, Singapore (OFIRG23jan-0057 to LTS)

The Ministry of Education, Singapore (MOE-T2EP30220-0012 and MOE-T2EP30224-0012 to LTS; MOE-T2EP30223-0002 and MOE-T2EP10124-0001 to Y.Q.)

The National Research Foundation Singapore (NRF-F-CRP-2024-0002 to LTS) National Institute of Health Grants (5R35GM150669 to JFK)

Smith Family Foundation (Grant for Excellence in Biomedical Research to JFK)

UMass Chan (Startup funds and BMB Excellence Award Grant to JFK)

This material by ENIGMA- Ecosystems and Networks Integrated with Genes and Molecular Assemblies (http://enigma.lbl.gov), a Science Focus Area Program at Lawrence Berkeley National Laboratory, is based upon work supported by the U.S. Department of Energy, Office of Science, Office of Biological & Environmental Research under contract number DE-AC02-05CH11231

## Author Contributions

Conceptualization: J. Z. and L-T. S.

Methodology: J. Z., Z. F., Y. L., Q. Y., and L.-T. S.

Investigation: J. Z., Z. F., Y. L., Q. Y., J.-F. K., and L.-T. S.

Visualization: J. Z., Z. F., Y. L., Q. Y., J.-F. K., and L.-T. S.

Supervision: Q. Y., J.-F. K., and L.-T. S.

Writing – Original draft: J. Z. and L-T. S.

Writing – review & editing: J. Z., Z. F., Y. L., Q. Y., J.-F. K., and L.-T. S.

## Data and Materials Availability

All data are available in the main text or the supplementary materials.

## REFERENCES

1. J. N. Weiser, D. M. Ferreira, J. C. Paton, *Streptococcus pneumoniae*: transmission, colonization and invasion. Nat Rev Microbiol 16, 355–367 (2018).

2. GBD 2019 Antimicrobial Resistance Collaborators, Global mortality associated with 33 bacterial pathogens in 2019: a systematic analysis for the Global Burden of Disease Study 2019. Lancet S0140-6736(22)02185–7 (2022). 10.1016/S0140-6736(22)02185-7.

3. R. G. Bender, et al., Global, regional, and national incidence and mortality burden of non-COVID-19 lower respiratory infections and aetiologies, 1990–2021: a systematic analysis from the Global Burden of Disease Study 2021. The Lancet Infectious Diseases 24, 974–1002 (2024).

4. B. Wahl, et al., Burden of *Streptococcus pneumoniae* and *Haemophilus influenzae* type b disease in children in the era of conjugate vaccines: global, regional, and national estimates for 2000-15. Lancet Glob Health 6, e744–e757 (2018).

5. M. Naghavi, et al., Global burden of bacterial antimicrobial resistance 1990–2021: a systematic analysis with forecasts to 2050. The Lancet 404, 1199–1226 (2024).

6. Active Bacterial Core Surveillance (ABCs) Report Emerging Infections Program Network=: *Streptococcus pneumoniae*, 2018. Available at: https://stacks.cdc.gov/view/cdc/109750 [Accessed 23 February 2024].

7. WHO publishes list of bacteria for which new antibiotics are urgently needed. Available at: https://www.who.int/news-room/detail/27-02-2017-who-publishes-list-of-bacteria-for-which-new-antibiotics-are-urgently-needed [Accessed 17 July 2020].

8. E. Tacconelli, et al., Discovery, research, and development of new antibiotics: the WHO priority list of antibiotic-resistant bacteria and tuberculosis. Lancet Infect Dis 18, 318–327 (2018).

9. R. Martinez-Vega, et al., Risk factor profiles and clinical outcomes for children and adults with pneumococcal infections in Singapore: A need to expand vaccination policy? PLOS ONE 14, e0220951 (2019).

10. P. Eng, et al., Role of pneumococcal vaccination in prevention of pneumococcal disease among adults in Singapore. Int J Gen Med 7, 179–191 (2014).

11. J. D. Grabenstein, K. P. Klugman, A century of pneumococcal vaccination research in humans. Clin Microbiol Infect 18 **Suppl 5**, 15–24 (2012).

12. F. A. Ganaie, et al., Update on the evolving landscape of pneumococcal capsule types: new discoveries and way forward. Clin Microbiol Rev 38, e0017524 (2025).

13. C. K.-R. Wong, Y.-Y. Chun, T. Su, L.-T. Sham, Leveraging the Capsular Polysaccharide Synthesis Pathway in *Streptococcus pneumoniae* as a Genetic Glycoengineering Platform. ACS Bio Med Chem Au 5, 342–349 (2025).

14. H. An, et al., Functional vulnerability of liver macrophages to capsules defines virulence of blood-borne bacteria. J Exp Med 219, e20212032 (2022).

15. X. Tian, et al., Natural antibodies to polysaccharide capsules enable Kupffer cells to capture invading bacteria in the liver sinusoids. J Exp Med 222, e20240735 (2024).

16. Y.-Y. Chun, et al., Influence of glycan structure on the colonization of *Streptococcus pneumoniae* on human respiratory epithelial cells. Proc Natl Acad Sci U S A 120, e2213584120 (2023).

17. T. Su, et al., Decoding capsule synthesis in *Streptococcus pneumoniae*. FEMS Microbiol Rev 45, fuaa067 (2021).

18. W.-Z. Chua, et al., High-throughput mutagenesis and cross-complementation experiments reveal substrate preference and critical residues of the capsule transporters in *Streptococcus pneumoniae*. mBio 12, e02615–21 (2021).

19. W.-Z. Chua, et al., Massively parallel barcode sequencing revealed the interchangeability of capsule transporters in *Streptococcus pneumoniae*. Sci Adv 11, eadr0162 (2025).

20. B. Xayarath, J. Yother, Mutations blocking side chain assembly, polymerization, or transport of a Wzy-dependent *Streptococcus pneumoniae* capsule are lethal in the absence of suppressor mutations and can affect polymer transfer to the cell wall. J Bacteriol 189, 3369–3381 (2007).

21. T. R. Larson, J. Yother, *Streptococcus pneumoniae* capsular polysaccharide is linked to peptidoglycan via a direct glycosidic bond to β-D-N-acetylglucosamine. Proc Natl Acad Sci U S A 114, 5695–5700 (2017).

22. R. Nakamoto, et al., The bacterial tyrosine kinase system CpsBCD governs the length of capsule polymers. Proc Natl Acad Sci U S A 118, e2103377118 (2021).

23. J. Yother, Capsules of *Streptococcus pneumoniae* and other bacteria: paradigms for polysaccharide biosynthesis and regulation. Annu Rev Microbiol 65, 563–81 (2011).

24. R. Nakamoto, et al., The divisome but not the elongasome organizes capsule synthesis in *Streptococcus pneumoniae*. Nat Commun 14, 3170 (2023).

25. J. Figueiredo, et al., Encapsulation of the septal cell wall protects *Streptococcus pneumoniae* from its major peptidoglycan hydrolase and host defenses. PLoS Pathog 18, e1010516 (2022).

26. M. X. Henriques, T. Rodrigues, M. Carido, L. Ferreira, S. R. Filipe, Synthesis of capsular polysaccharide at the division septum of *Streptococcus pneumoniae* is dependent on a bacterial tyrosine kinase. Mol Microbiol 82, 515–34 (2011).

27. J. Nourikyan, et al., Autophosphorylation of the bacterial tyrosine-kinase CpsD connects capsule synthesis with the cell cycle in *Streptococcus pneumoniae*. PLoS Genet 11, e1005518 (2015).

28. Y. Kawai, et al., A widespread family of bacterial cell wall assembly proteins. EMBO J 30, 4931–41 (2011).

29. K. Schaefer, L. M. Matano, Y. Qiao, D. Kahne, S. Walker, In vitro reconstitution demonstrates the cell wall ligase activity of LCP proteins. Nat Chem Biol 13, 396–401 (2017).

30. Y. G. Chan, H. K. Kim, O. Schneewind, D. Missiakas, The capsular polysaccharide of *Staphylococcus aureus* is attached to peptidoglycan by the LytR-CpsA-Psr (LCP) family of enzymes. J Biol Chem 289, 15680–90 (2014).

31. Y. G. Chan, M. B. Frankel, V. Dengler, O. Schneewind, D. Missiakas, *Staphylococcus aureus* mutants lacking the LytR-CpsA-Psr family of enzymes release cell wall teichoic acids into the extracellular medium. J Bacteriol 195, 4650–9 (2013).

32. J. Hübscher, L. Lüthy, B. Berger-Bächi, P. Stutzmann Meier, Phylogenetic distribution and membrane topology of the LytR-CpsA-Psr protein family. BMC Genomics 9, 617 (2008).

33. K. Schaefer, T. W. Owens, D. Kahne, S. Walker, Substrate preferences establish the order of cell wall assembly in *Staphylococcus aureus*. J Am Chem Soc 140, 2442–2445 (2018).

34. N. Hess, et al., Lipoteichoic acid deficiency permits normal growth but impairs virulence of *Streptococcus pneumoniae*. Nat Commun 8, 2093 (2017).

35. W. Fischer, T. Behr, R. Hartmann, J. Peter-Katalinić, H. Egge, Teichoic acid and lipoteichoic acid of *Streptococcus pneumoniae* possess identical chain structures. A reinvestigation of teichoid acid (C polysaccharide). Eur J Biochem 215, 851–857 (1993).

36. A. Eberhardt, et al., Attachment of capsular polysaccharide to the cell wall in *Streptococcus pneumoniae*. Microb Drug Resist 18, 240–255 (2012).

37. W. Ye, et al., Pneumococcal LytR protein is required for the surface attachment of both capsular polysaccharide and teichoic acids: essential for pneumococcal virulence. Front Microbiol 9, 1199 (2018).

38. J. K. Morona, D. C. Miller, R. Morona, J. C. Paton, The effect that mutations in the conserved capsular polysaccharide biosynthesis genes *cpsA*, *cpsB*, and *cpsD* Have on Virulence of *Streptococcus pneumoniae*. J Infect Dis 189, 1905–1913 (2004).

39. M. H. Bender, R. T. Cartee, J. Yother, Positive correlation between tyrosine phosphorylation of CpsD and capsular polysaccharide production in *Streptococcus pneumoniae*. J Bacteriol 185, 6057–66 (2003).

40. J. K. Morona, J. C. Paton, D. C. Miller, R. Morona, Tyrosine phosphorylation of CpsD negatively regulates capsular polysaccharide biosynthesis in *Streptococcus pneumoniae*. Mol Microbiol 35, 1431–1442 (2000).

41. O. Johnsborg, L. S. Håvarstein, Pneumococcal LytR, a protein from the LytR-CpsA-Psr family, is essential for normal septum formation in *Streptococcus pneumoniae*. J Bacteriol 191, 5859–5864 (2009).

42. L. Cuthbertson, I. L. Mainprize, J. H. Naismith, C. Whitfield, Pivotal roles of the outer membrane polysaccharide export and polysaccharide copolymerase protein families in export of extracellular polysaccharides in gram-negative bacteria. Microbiol Mol Biol Rev 73, 155–177 (2009).

43. B. Wiseman, R. G. Nitharwal, G. Widmalm, M. Högbom, Structure of a full-length bacterial polysaccharide co-polymerase. Nat Commun 12, 369 (2021).

44. S. T. Islam, J. S. Lam, Synthesis of bacterial polysaccharides via the Wzx/Wzy-dependent pathway. Can J Microbiol 60, 697–716 (2014).

45. Y.-H. Chen, et al., Facile manipulation of protein localization in fission yeast through binding of GFP-binding protein to GFP. J Cell Sci 130, 1003–1015 (2017).

46. K. M. Wetmore, et al., Rapid quantification of mutant fitness in diverse bacteria by sequencing randomly bar-coded transposons. mBio 6, e00306–15 (2015).

47. J. J. Zik, et al., Dual transposon sequencing profiles the genetic interaction landscape in bacteria. Science 389, eadt7685 (2025).

48. V. D. H. Ding, et al., Purification and characterization of recombinant human farnesyl diphosphate synthase expressed in *Escherichia coli*. Biochem J 275, 61–65 (1991).

49. S. Heuston, M. Begley, C. G. M. Gahan, C. Hill, Isoprenoid biosynthesis in bacterial pathogens. Microbiology 158, 1389–1401 (2012).

50. J. Flores-Kim, G. S. Dobihal, A. Fenton, D. Z. Rudner, T. G. Bernhardt, A switch in surface polymer biogenesis triggers growth-phase-dependent and antibiotic-induced bacteriolysis. Elife 8, e44912 (2019).

51. S. M. Barendt, et al., Influences of capsule on cell shape and chain formation of wild-type and *pcsB* mutants of serotype 2 *Streptococcus pneumoniae*. J Bacteriol 191, 3024–3040 (2009).

52. L. Kong, P. Zhang, P. Setlow, Y.-Q. Li, Characterization of bacterial spore germination using integrated phase contrast microscopy, Raman spectroscopy, and optical tweezers. Anal Chem 82, 3840–3847 (2010).

53. M. Nguyen, et al., Teichoic acids in the periplasm and cell envelope of *Streptococcus pneumoniae*. eLife 14, e105132 (2025).

54. M. E. Winkler, J. A. Hoch, Essentiality, bypass, and targeting of the YycFG (VicRK) two-component regulatory system in gram-positive bacteria. J Bacteriol 190, 2645–8 (2008).

55. W. L. Ng, M. E. Winkler, Singular structures and operon organizations of essential two-component systems in species of *Streptococcus*. Microbiology 150, 3096–8 (2004).

56. S. Dubrac, P. Bisicchia, K. M. Devine, T. Msadek, A matter of life and death: cell wall homeostasis and the WalKR (YycGF) essential signal transduction pathway. Mol Microbiol 70, 1307–1322 (2008).

57. S. Dubrac, T. Msadek, Tearing down the wall: peptidoglycan metabolism and the WalK/WalR (YycG/YycF) essential two-component system. Adv Exp Med Biol 631, 214–228 (2008).

58. G. S. Dobihal, Y. R. Brunet, J. Flores-Kim, D. Z. Rudner, Homeostatic control of cell wall hydrolysis by the WalRK two-component signaling pathway in *Bacillus subtilis*. Elife 8, e52088 (2019).

59. S. Dubrac, I. G. Boneca, O. Poupel, T. Msadek, New insights into the WalK/WalR (YycG/YycF) essential signal transduction pathway reveal a major role in controlling cell wall metabolism and biofilm formation in *Staphylococcus aureus*. J Bacteriol 189, 8257–8269 (2007).

60. W. L. Ng, H. C. Tsui, M. E. Winkler, Regulation of the *pspA* virulence factor and essential *pcsB* murein biosynthetic genes by the phosphorylated VicR (YycF) response regulator in *Streptococcus pneumoniae*. J Bacteriol 187, 7444–59 (2005).

61. R. Lange, et al., Domain organization and molecular characterization of 13 two-component systems identified by genome sequencing of *Streptococcus pneumoniae*. Gene 237, 223–234 (1999).

62. W.-L. Ng, et al., Constitutive expression of PcsB suppresses the requirement for the essential VicR (YycF) response regulator in *Streptococcus pneumoniae* R6. Mol Microbiol 50, 1647–1663 (2003).

63. J. P. Throup, et al., A genomic analysis of two-component signal transduction in *Streptococcus pneumoniae*. Mol Microbiol 35, 566–576 (2000).

64. D. B. Senadheera, et al., Regulation of bacteriocin production and cell death by the VicRK signaling system in *Streptococcus mutans*. J Bacteriol 194, 1307–1316 (2012).

65. M. Liu, et al., Defects in ex vivo and in vivo growth and sensitivity to osmotic stress of group A *Streptococcus* caused by interruption of response regulator gene *vicR*. Microbiology (Reading) 152, 967–978 (2006).

66. G. S. Dobihal, J. Flores-Kim, I. J. Roney, X. Wang, D. Z. Rudner, The WalR-WalK signaling pathway modulates the activities of both cwlo and lyte through control of the peptidoglycan deacetylase PdaC in *Bacillus subtilis*. J Bacteriol 204, e0053321 (2022).

67. K. J. Wayne, et al., Localization and cellular amounts of the WalRKJ (VicRKX) two-component regulatory system proteins in serotype 2 *Streptococcus pneumoniae*. J Bacteriol 192, 4388–4394 (2010).

68. N. S. Briggs, et al., Regulation of the essential peptidoglycan hydrolytic complex FtsEX-PcsB during *Streptococcus pneumoniae* cell division. bioRxiv 2025.05.23.655736 (2025). 10.1101/2025.05.23.655736.

69. P. García, M. P. González, E. García, R. López, J. L. García, LytB, a novel pneumococcal murein hydrolase essential for cell separation. Mol Microbiol 31, 1275–1281 (1999).

70. V. Miguel-Ruano, et al., Characterization of VldE (Spr1875), a pneumococcal two-state L,D-endopeptidase with a four-zinc cluster in the active site. ACS Catal 14, 18786–18798 (2024).

71. J. Zhang, et al., Inactivation of transcriptional regulator FabT influences colony phase variation of *Streptococcus pneumoniae*. mBio 12, e0130421 (2021).

72. J. Flores-Kim, G. S. Dobihal, T. G. Bernhardt, D. Z. Rudner, WhyD tailors surface polymers to prevent premature bacteriolysis and direct cell elongation in *Streptococcus pneumoniae*. Elife 11, e76392 (2022).

73. W.-L. Ng, K. M. Kazmierczak, M. E. Winkler, Defective cell wall synthesis in *Streptococcus pneumoniae* R6 depleted for the essential PcsB putative murein hydrolase or the VicR (YycF) response regulator. Mol Microbiol 53, 1161–1175 (2004).

74. J. D. Cohen, Evidence that glycopolymer transferases promote peptidoglycan hydrolysis in *Bacillus subtilis*. bioRxiv 2025.02.26.640348 (2025). 10.1101/2025.02.26.640348.

75. D. Denapaite, R. Bruckner, R. Hakenbeck, W. Vollmer, Biosynthesis of teichoic acids in *Streptococcus pneumoniae* and closely related species: lessons from genomes. Microb Drug Resist 18, 344–58 (2012).

76. W. P. Robins, S. M. Faruque, J. J. Mekalanos, Coupling mutagenesis and parallel deep sequencing to probe essential residues in a genome or gene. Proc Natl Acad Sci U S A 110, E848–E857 (2013).

77. Y. Li, C. M. Thompson, M. Lipsitch, A modified *Janus* cassette (Sweet *Janus*) to improve allelic replacement efficiency by high-stringency negative selection in *Streptococcus pneumoniae*. PLoS One 9, e100510 (2014).

78. C. Wagner, et al., Genetic analysis and functional characterization of the *Streptococcus pneumoniae* vic operon. Infect Immun 70, 6121–6128 (2002).

79. J. P. Lisher, et al., Biological and chemical adaptation to endogenous hydrogen peroxide production in *Streptococcus pneumoniae* D39. mSphere 2, e00291–16 (2017).

80. K. A. Geno, J. R. Hauser, K. Gupta, J. Yother, *Streptococcus pneumoniae* phosphotyrosine phosphatase CpsB and alterations in capsule production resulting from changes in oxygen availability. J Bacteriol 196, 1992–2003 (2014).

81. S. M. Carvalho, et al., Pyruvate Oxidase Influences the Sugar Utilization Pattern and Capsule Production in *Streptococcus pneumoniae*. PLOS ONE 8, e68277 (2013).

82. H. Yesilkaya, V. F. Andisi, P. W. Andrew, J. J. E. Bijlsma, *Streptococcus pneumoniae* and reactive oxygen species: an unusual approach to living with radicals. Trends Microbiol 21, 187–195 (2013).

83. D. G. Glanville, et al., Pneumococcal capsule expression is controlled through a conserved, distal cis-regulatory element during infection. PLoS Pathog 19, e1011035 (2023).

84. G. A. Stamsås, D. Straume, Z. Salehian, L. S. Håvarstein, Evidence that pneumococcal WalK is regulated by StkP through protein-protein interaction. Microbiology (Reading) 163, 383–399 (2017).

85. V. Dengler, et al., Deletion of hypothetical wall teichoic acid ligases in *Staphylococcus aureus* activates the cell wall stress response. FEMS Microbiol Lett 333, 109–120 (2012).

86. C. Stefanović, F. F. Hager, C. Schäffer, LytR-CpsA-Psr Glycopolymer Transferases: Essential Bricks in Gram-Positive Bacterial Cell Wall Assembly. Int J Mol Sci 22, 908 (2021).

87. T. Sandalova, et al., Crystallographic and NMR studies of *Streptococcus pneumoniae* LCP protein PsrSp indicate the importance of dynamics in four long loops for ligand specificity. Crystals 14, 1094 (2024).

88. F. K. K. Li, et al., Crystallographic analysis of *Staphylococcus aureus* LcpA, the primary wall teichoic acid ligase. J Biol Chem 295, 2629–2639 (2020).

89. A. Rajaei, H. M. Rowe, M. N. Neely, The LCP Family Protein, Psr, Is Required for Cell Wall Integrity and Virulence in *Streptococcus agalactiae*. Microorganisms 10, 217 (2022).

90. M. Baumgart, K. Schubert, M. Bramkamp, J. Frunzke, Impact of LytR-CpsA-Psr Proteins on Cell Wall Biosynthesis in *Corynebacterium glutamicum*. J Bacteriol 198, 3045–3059 (2016).

91. W. Vollmer, Structural variation in the glycan strands of bacterial peptidoglycan. FEMS Microbiol Rev 32, 287–306 (2008).

92. H. Takada, et al., Essentiality of WalRK for growth in *Bacillus subtilis* and its role during heat stress. Microbiology (Reading) 164, 670–684 (2018).

93. F. E. Jacobsen, K. M. Kazmierczak, J. P. Lisher, M. E. Winkler, D. P. Giedroc, Interplay between manganese and zinc homeostasis in the human pathogen *Streptococcus pneumoniae*. Metallomics 3, 38–41 (2011).

94. J. E. Martin, J. P. Lisher, M. E. Winkler, D. P. Giedroc, Perturbation of manganese metabolism disrupts cell division in *Streptococcus pneumoniae*. Mol Microbiol 104, 334–348 (2017).

95. X. Liu, et al., High-throughput CRISPRi phenotyping identifies new essential genes in *Streptococcus pneumoniae*. Mol Syst Biol 13, 931 (2017).

96. D. G. Gibson, et al., Enzymatic assembly of DNA molecules up to several hundred kilobases. Nat Methods 6, 343–345 (2009).

97. R. Junges, et al., Markerless Genome Editing in Competent *Streptococci*. Methods Mol Biol 1537, 233–247 (2017).

98. T. Su, et al., Rewiring the pneumococcal capsule pathway for investigating glycosyltransferase specificity and genetic glycoengineering. Sci Adv 9, eadi8157 (2023).

99. C. K. Sung, H. Li, J. P. Claverys, D. A. Morrison, An *rpsL* cassette, *janus*, for gene replacement through negative selection in *Streptococcus pneumoniae*. Appl Environ Microbiol 67, 5190–5196 (2001).

100. K. J. Livak, T. D. Schmittgen, Analysis of relative gene expression data using real-time quantitative PCR and the 2(-Delta Delta C(T)) Method. Methods 25, 402–408 (2001).

101. J. Schindelin, et al., Fiji: an open-source platform for biological-image analysis. Nat Methods 9, 676–682 (2012).

