## Supplementary figures and images for "The WalRK two-component system in *Streptococcus pneumoniae* ensures robustness of secondary wall polymer attachment"

### Fig. S1

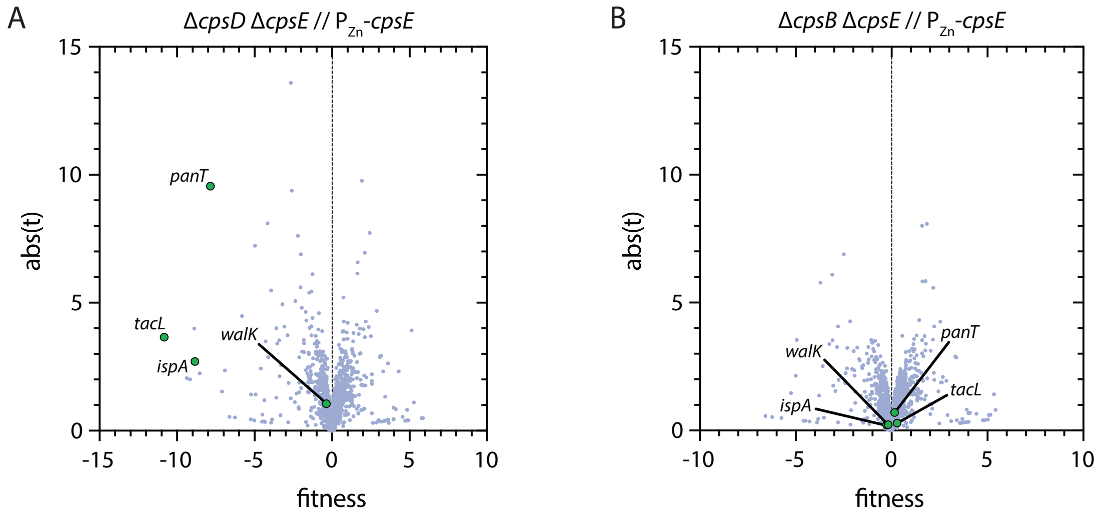

### Fig. S2

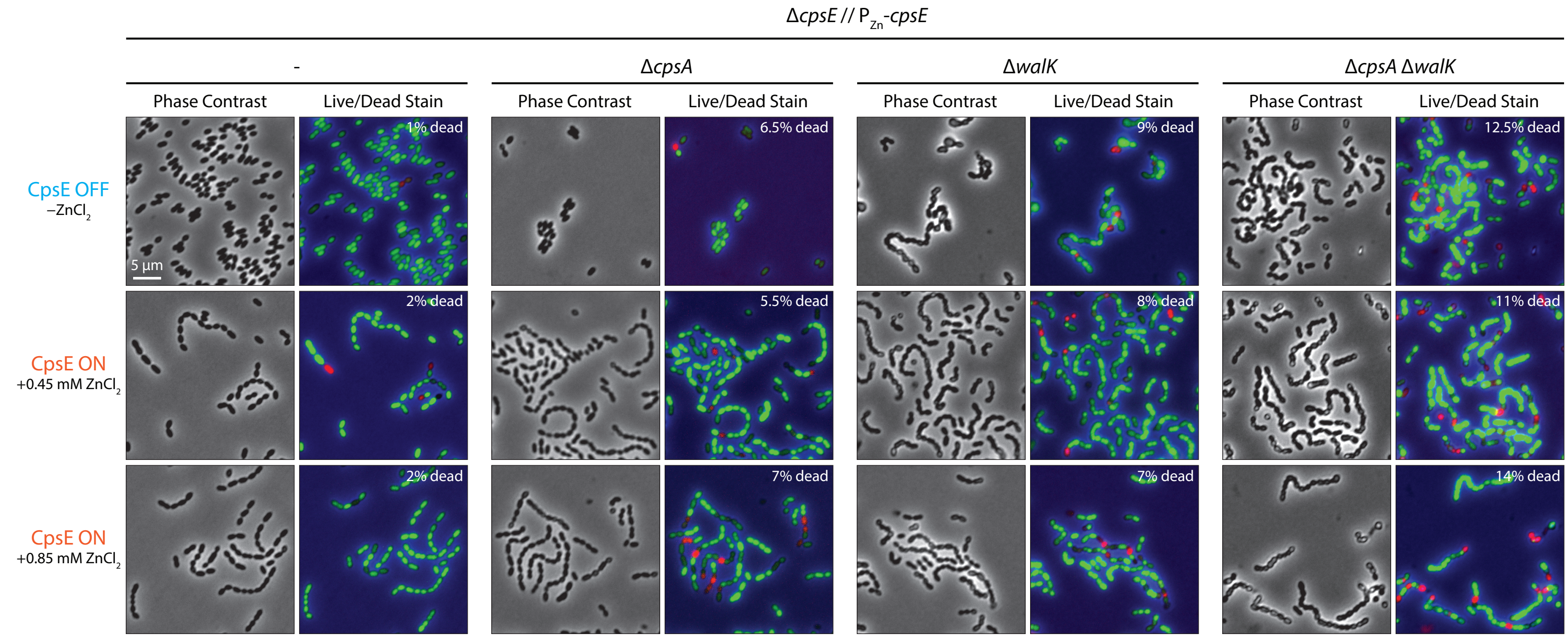

### Fig. S3

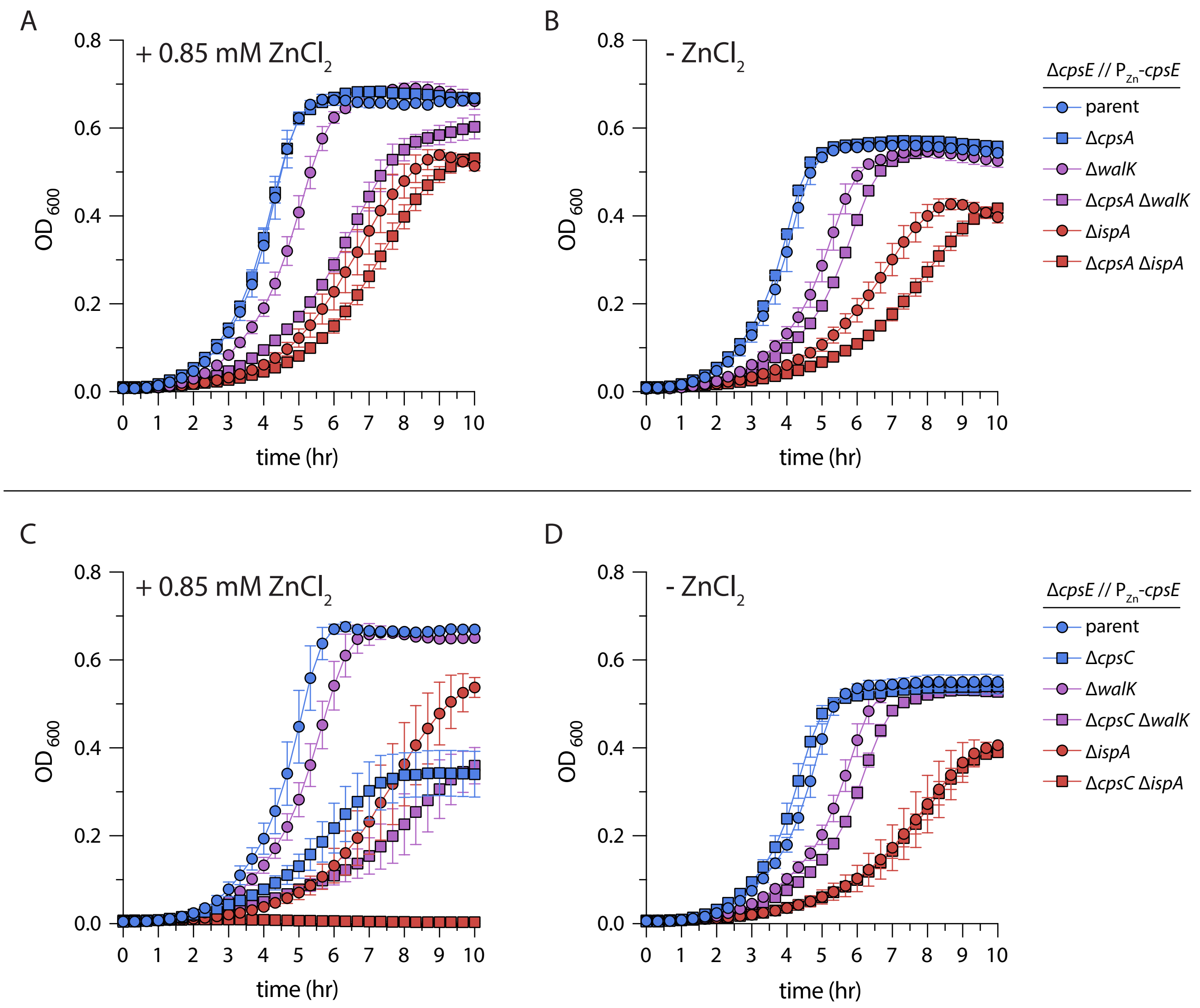

### Fig. S4

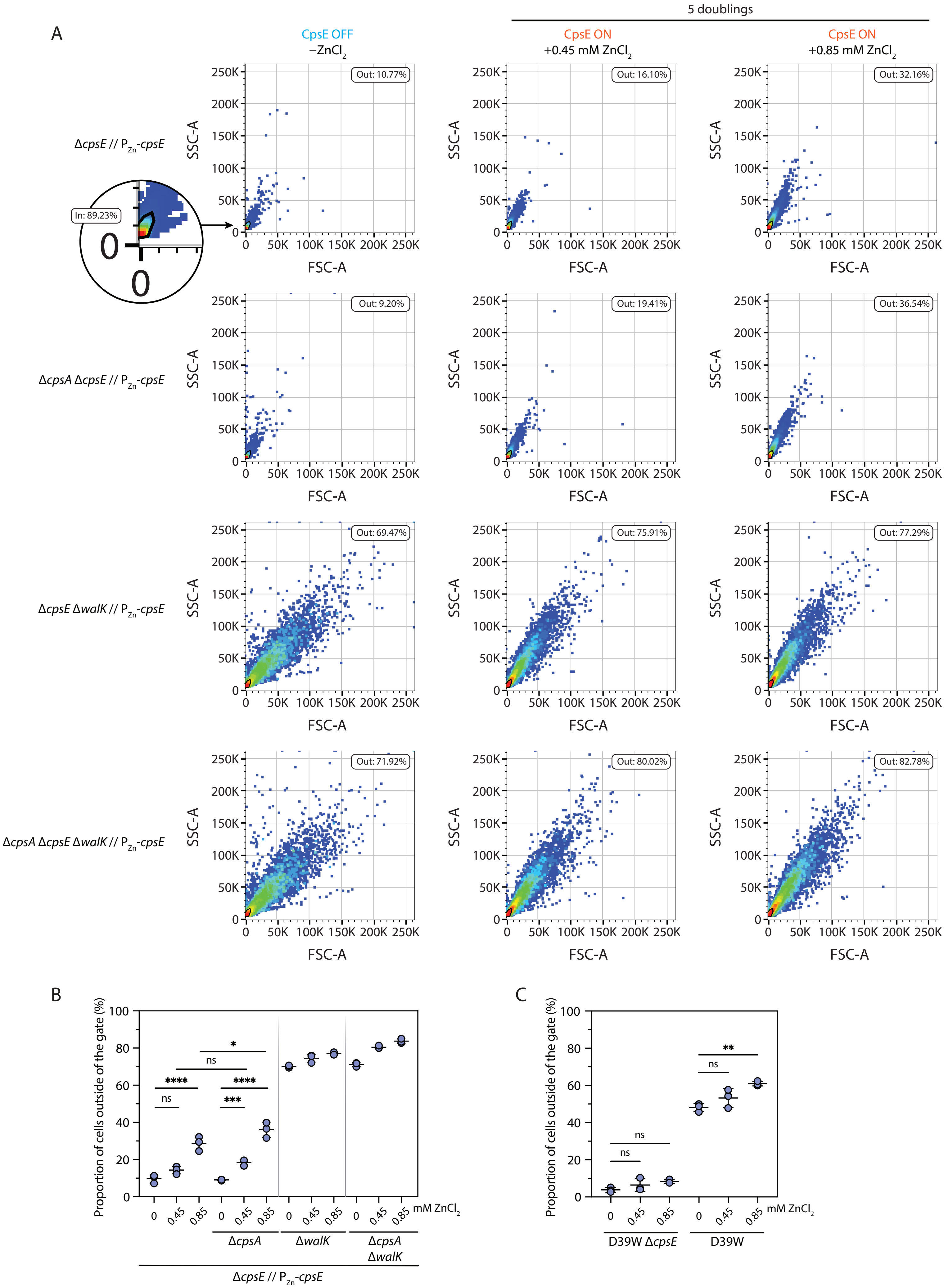

### Fig. S5

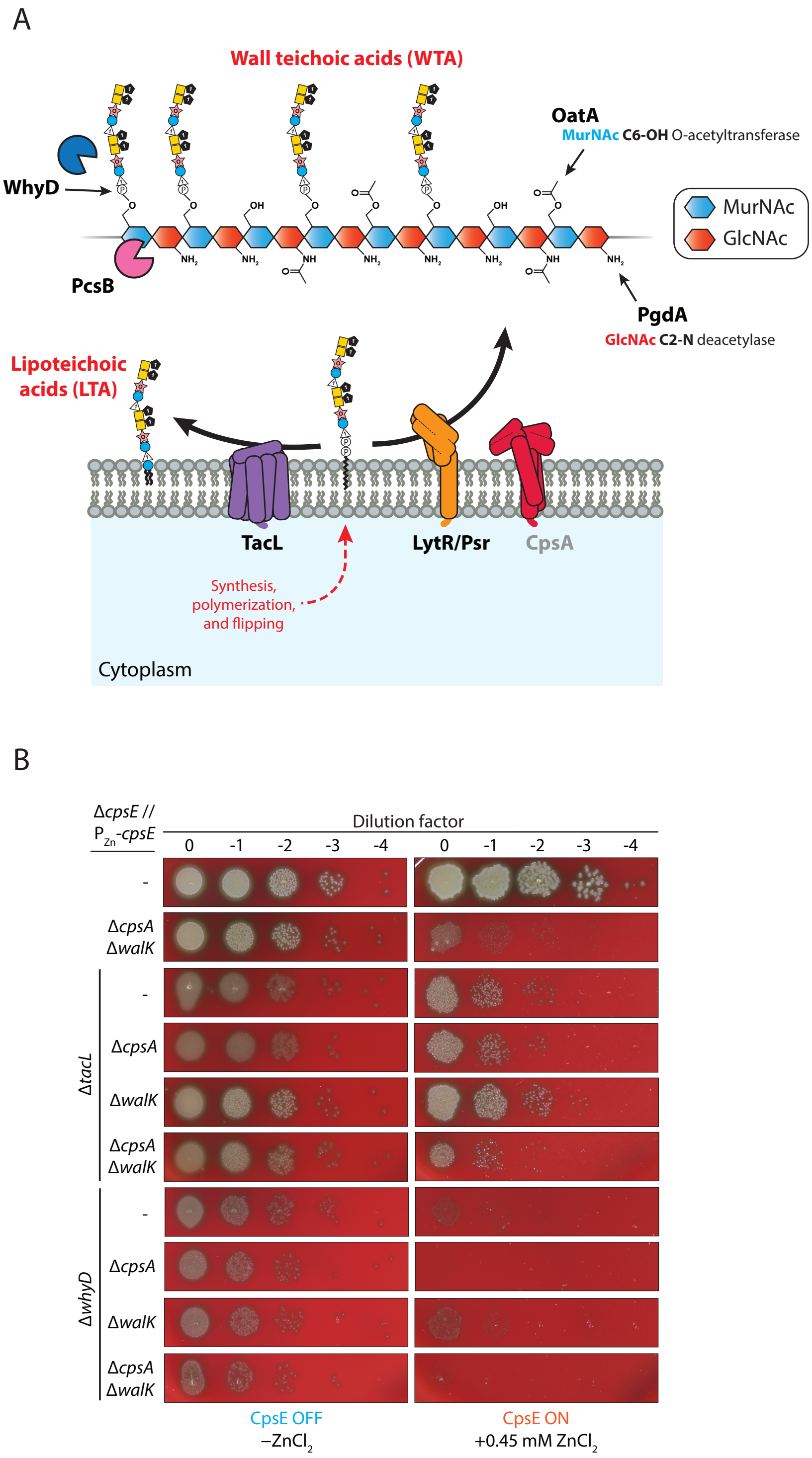

### Fig. S6

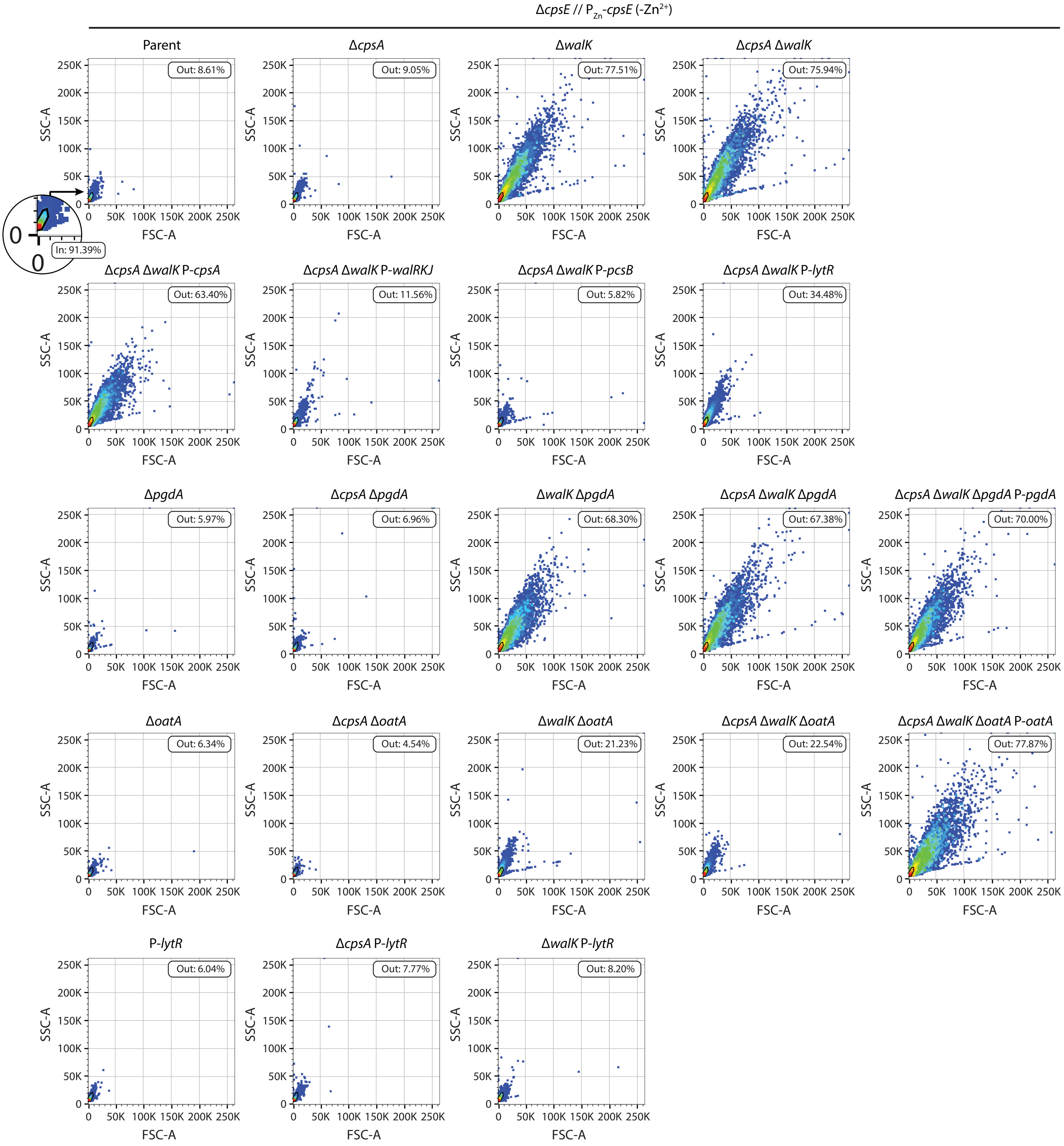

### Fig. S7

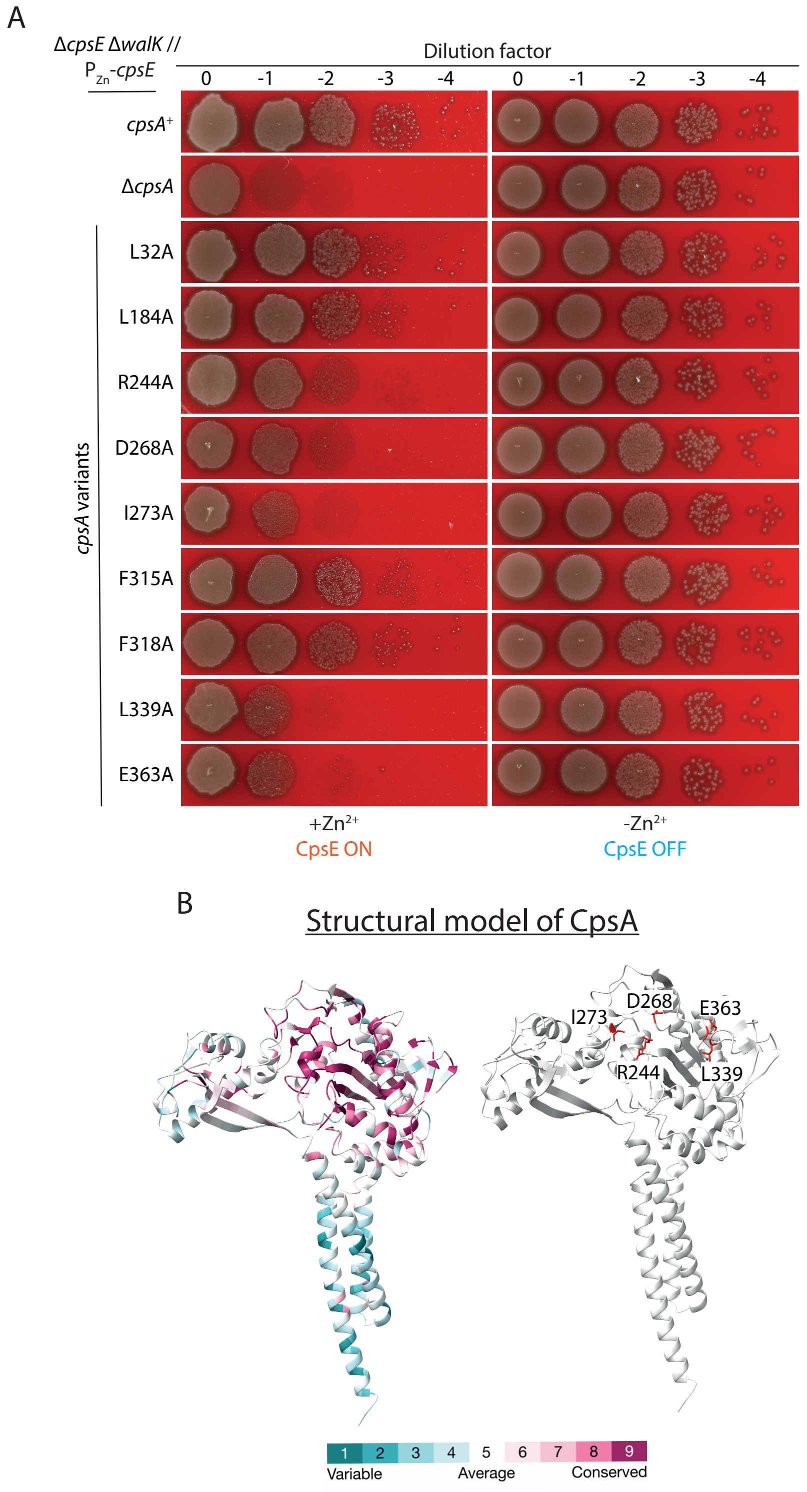

### Fig. S8

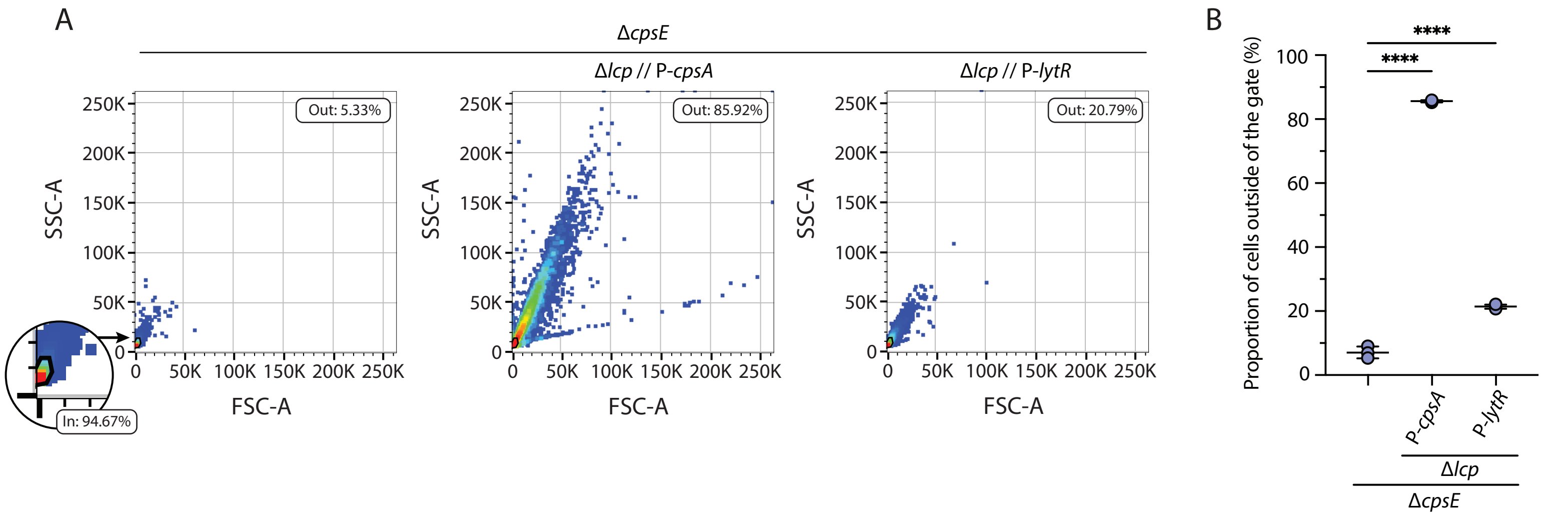
